# A systematic benchmark of bioinformatics methods for single-cell and spatial RNA-seq Nanopore long-read data

**DOI:** 10.1101/2025.07.21.665920

**Authors:** Ali Hamraoui, Audrey Onfroy, Catherine Senamaud-Beaufort, Fanny Coulpier, Sophie Lemoine, Laurent Jourdren, Morgane Thomas-Chollier

## Abstract

Alternative splicing plays a crucial role in transcriptomic complexity, yet remains difficult to resolve at the single-cell level due to the limitations of short-read technologies. Coupling single-cell with long-read sequencing offers full-length transcript coverage, enabling more accurate isoform detection. Multiple specialized computational tools tailored for single-cell and spatial long-read transcriptomics have been developed, with diverse strategies. To compare the effectiveness of these approaches, we generated paired short-read and Nanopore long-read single-cell datasets, tailored for benchmarking bioinformatics tools. We evaluated ten state-of-the-art methods, spanning four analytical dimensions: barcodes and UMI detection, demultiplexing and UMI clustering, gene-level expression profiling, and isoform detection and quantification. Using real and simulated datasets across different protocols, sequencing depths and chemistries, we assessed the accuracy, robustness, and scalability of each tool. Our results revealed method-specific trade-offs, and highlight the importance of sequencing quality and UMI correction strategies. This benchmark provides a practical resource for optimizing isoform analysis and accurate gene expression profiling in single-cell and spatial transcriptomics using long-read sequencing. Our benchmarking workflow is designed to be reusable, thereby enabling method developers to compare their own approaches against the set of reference methods evaluated in this work.

## Introduction

Alternative splicing significantly contributes to transcriptome complexity and has critical implications for cellular functions. While identifying isoforms can be achieved using bulk RNA-sequencing, this approach relies on RNA pooled from thousands of cells, resulting in averaged transcriptome information that ignores cell heterogeneity (1). Recent advancements in single-cell isolation and capture techniques, such as droplet-based methods (2, 3), and in situ-capture-based methods (4, 5) have enabled high-throughput quantification of gene expression at both single-cell resolution and in spatially resolved contexts.

However, high-throughput full-length RNA isoform identification and quantification remain challenging at the single-cell level. Typical droplet-based single-cell protocols, sequenced in short reads, capture only limited sequence information. This information is typically restricted to one end of the transcript (e.g. 10X Genomics 3’ Chemistry) and does not span exon-exon junctions across full-length isoforms. As a result, the comprehensive detection and quantification of alternative isoforms remains challenging (6, 7).

Plate-based protocols (e.g. smart-seq) are used as an alternative, for their high sensitivity and ability to capture entire transcripts (8). Nevertheless, they still suffer from intrinsic limitations of short-read technologies, leading to isoform reconstruction errors caused by ambiguous read-to-isoform mapping and 5’ coverage bias (9, 10). Furthermore, their limited throughput—generally around 400 cells per experiment—and relatively high cost per cell, reduce their suitability for large-scale studies (8, 11, 12).

Long-read sequencing technologies, including Oxford Nanopore Technology (ONT) and Pacific Biosciences (PacBio), overcome these limitations by enabling the unambiguous identification of complete exon structures (13). These sequencing platforms were previously limited by high sequencing error rates and/or low throughput. For single-cell approaches, this hampered the accurate detection of barcodes and unique molecular identifiers (UMIs), thus limiting the representation of the whole diversity of captured RNA transcripts (14). However, improvements of platforms and library preparation methods over the past five years have enhanced accuracy and throughput, with tens of millions of reads generated in a single experiment (15).

Beside technological development, several computational methods have been developed to specifically address bioinformatics challenges associated with processing long-read single-cell RNA-sequencing (scRNA-seq) or spatial data. Tools such as Sicelore (16), Snuupy (17), ScNapBar (18) and scTagger (19) focus on barcodes and UMI assignments by integrating paired short-read data. In contrast, approaches like Sicelore 2.1 (16), wf-single-cell (Oxford Nanopore Technology), scNanoGPS (20), Longcell (21) and Scywalker (22) perform barcodes demultiplexing and UMI clustering using only long-read data.

Similarly, methods such as FLAMES (23) and Bambu (24) manage multiple stages of the analysis pipeline, ranging from read alignment and UMI deduplication to isoform quantification and discovery, while relying on external tools like BLAZE (25) and flexiplex (26) for barcodes and UMI identification. Furthermore, tools such as Isosceles (27) and IsoQuant (28) build upon established demultiplexing and UMI correction strategies, such as those implemented in Sicelore 2.1 and wf-single-cell—to enhance transcript-level discovery and quantification.

Most recently, the nf-core community released a dedicated pipeline within the nf-core framework (29, 30), which integrates components such as BLAZE and IsoQuant for gene- and transcript-level quantification (31). A comprehensive overview of these methodological developments was recently provided by Gupta et al. (32) and Kumari et al. (33).

Yet, given the increasing applications of single-cell and spatial long-read sequencing in exploring transcriptome heterogeneity, evaluating and comparing these computational methods becomes crucial. Notably, bioinformatics software publications often present overly optimistic self-assessments (34, 35). Thus, independent external benchmarking is essential to address these biases effectively (36, 37). Additionally, accurate isoform detection is one of the key advantages of single-cell long-read sequencing technologies. While several isoform-level methods such as FLAMES, Bambu, and StringTie (integrated into wf-single-cell) have been evaluated extensively in the context of bulk RNA-seq (38, 39), their performance on single-cell long-read data remains less explored.

In this study, we systematically benchmarked state-of-the-art computational tools for single-cell and spatial long-read transcriptomics across four essential analytical dimensions. First, we assessed the number of detected UMIs and genes by comparing long-read approaches against short-read references. Second, we evaluated the barcodes identification and UMI correction and deduplication strategies, critical for accurate molecular identification and quantification. Third, we investigated the impact of these methods on transcriptomic profiling and cell annotation at the gene level. Lastly, we examined the performance of isoform detection, quantification, and discovery approaches. This comprehensive and independent benchmarking provides crucial insights for selecting optimal bioinformatics methods tailored to detect isoforms at the single-cell level.

## Results

### Overview of the study

#### Benchmarking datasets

This benchmark was performed using 20 long-read Nanopore datasets : six novel scRNA-seq datasets generated with 10X Genomics technology, two publicly available spatial Visium RNA-seq dataset, and simulated scRNA-seq data (Figure 1a and Table S1). The scRNA-seq data were newly-generated from two samples of malignant peripheral nerve sheath tumors (MPNSTs), designated as MPNST1 and MPNST2, developed by a mouse model of Neurofibromatosis Type 1 (40). First, we performed short-read sequencing using the Illumina platform, which yielded 491 and 401 million reads respectively, with a mean of 107,700 and 38,900 reads per cell, approximately 4,500 and 10,000 single cells, and a total of *~* 44 million and *~* 134 million UMIs associated with genes, achieving high sequencing saturations of 83% and 81% for MPNST1 and MPNST2 respectively. We then sequenced the same MPNST1 samples with two long-read ONT platforms to assess the impact of sequencing depth on methods performances. The MinION platform generated 11 million long reads, while the PromethION platform produced 151 million long reads, corresponding to sequencing saturations of 4% and 35%, respectively. Similarly, we sequenced the same MPNST2 samples solely on the PromethION platform, but following two distinct library preparation protocols: ScNaUmi-seq (16) and single-cell long-read ONT protocol (methods), which produced 130 million reads and 108 million reads, respectively. These two datasets allowed us to evaluate a possible impact of library preparation protocols on methods performances.

**Fig. 1.**
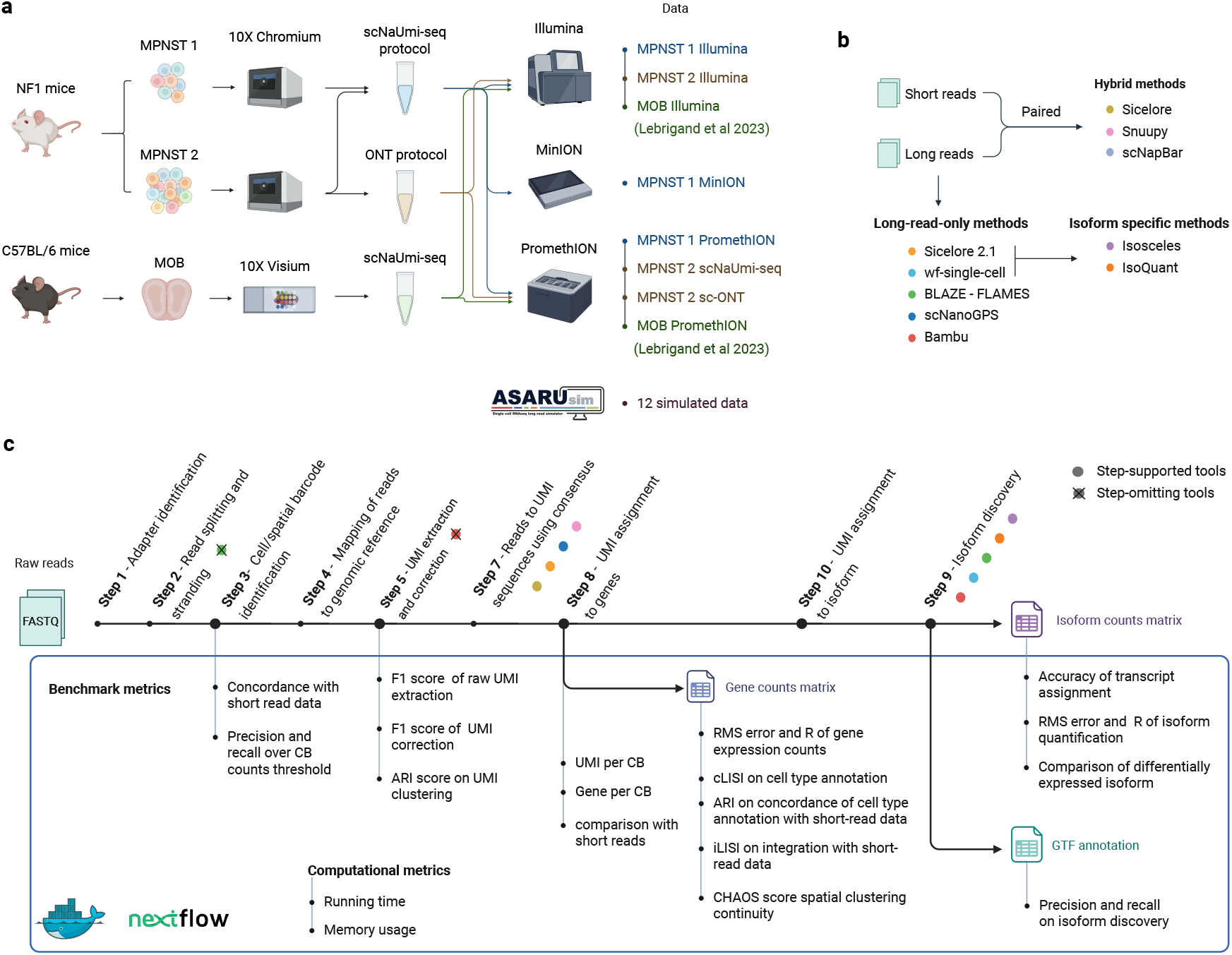
Overview of the benchmark —. **(a)** This study was performed with a total of 20 datasets on various cell types and sample sizes. Six were newly-generated (MPNST, 10X Genomics) covering two library preparation protocols and 3 sequencing platforms. Two were from a public dataset (MOB, Visium), and 12 were simulated with the dedicated tool AsaruSim. **(b)** 10 single-cell and spatial long-read RNA-seq methods were evaluated, organised into 3 categories : hybrid methods, long-read-only methods, and isoform-specific methods. **(c)** Data processing steps evaluated in this benchmark along with associated metrics. The preprocessing steps are organized chronologically, from raw FASTQ files to the generation of gene- and isoform-level count matrices. The step is supported by all methods, unless specified differently as follows : dots indicate the few methods supporting a particular step; crossed dots indicate the few methods that do not support a given step, with colors corresponding to the tools presented in panel b. The entire benchmark is implemented as a reusable workflow in Nextflow and executed within Docker containers.

Regarding the spatial long-read data, we downloaded a publicly available dataset, including FASTQ files and gene count matrix of the corresponding short-read data. This dataset comprises approximately 900 spatial spots from the olfactory bulb region of the mouse brain, generated using the Visium Spatial Gene Expression platform, and sequenced on ONT PromethION (41).

To assess tool performance against a well-defined ground truth, we generated various simulated scRNA-seq long-read datasets using the simulation framework AsaruSim (42) (Table S1).

Altogether, these datasets constitute a valuable resource for benchmarking long-read single-cell transcriptomics methods. By varying both sequencing platforms (MinION vs. PromethION), library preparation protocols (ScNaUmi-seq vs. ONT), R9.4.1 and R10.4.1 chemistry through simulations, they allow for a direct evaluation of the respective contributions of sequencing depth and protocol design to method performance.

#### Selected bioinformatics tools and their specificities

The first key challenge in scRNA-seq Nanopore data analysis lies in the accurate identification of barcodes and UMI sequences. We selected the popular tools through an extensive and systematic review conducted between 2022 and 2025, aiming to be as comprehensive as possible. Tools not included were either outdated, relying on custom tooling or ad hoc workflows, or lacked sufficient documentation. In brief, the ten computational tools can be grouped in three categories: hybrid, long-read-only and isoform-specific methods (Figure 1b-c, Table 2).

Hybrid methods, represented by Sicelore, Snuupy and scNapBar employ a hybrid sequencing approach in which barcodes and UMI assignment procedure is guided by short-read data. Sicelore extracts barcodes and UMI from short stretches of the long read between a valid adapter sequence and a threshold number of poly-A. Snuupy and scNapBar are developed as an improvement of Sicelore to either remove the dependency of the algorithm on the poly-A tail or the depth of short-read sequencing, respectively.

Conversely, long-read-only methods such as FLAMES, Sicelore 2.1, wf-single-cell, scNanoGPS, and Bambu, adopt long-read specific approaches, correcting barcodes and UMI without the use of paired short-read data. The long-read-only tools differ at three levels. Firstly, regarding barcode validation, FLAMES requires the cell-associated barcodes list as input, which can be generated by BLAZE or derived from a shortlist obtained from short-read sequencing data. Sicelore 2.1, wf-single-cell and Bambu (which relies on flexiplex) use an inclusion list (e.g. the *~*3 million unique sequences for 10X Genomics v3 chemistry) and retain the most highly represented barcodes within a defined count threshold. scNanoGPS applies the iCARLO algorithm (20) to identify and correct erroneous barcodes, focusing particularly on the most abundant sequences. Secondly, tools differ by their assignment of UMI sequences to reads. FLAMES, Sicelore 2.1, and scNanoGPS cluster or merge UMIs mapped to the same gene or genomic interval using a Levenshtein distance threshold. In contrast, wf-single-cell employs the directional graph approach from UMI-tools (43) to perform UMI clustering, while Bambu directly merges raw UMI sequences. Finally, regarding the selection of representative reads within each UMI cluster, FLAMES and Bambu retain the longest read. wf-single-cell selects a representative read—typically the one with the highest read count or the most “central” UMI—using the default strategy of UMI-tools. In contrast, Sicelore 2.1 and scNanoGPS generate a consensus sequence across all reads in the cluster (Table S2).

Isoform-specific methods, such as IsoQuant and Isosceles, use existing preprocessing methods (e.g. Sicelore and wf-single-cell) for barcodes and UMI detection, and performs reference-guided de novo detection, quantification, and downstream analysis of isoforms at single-cell and pseudo-bulk levels.

#### Benchmark and workflow

A comprehensive benchmark was conducted by applying a wide range of metrics to evaluate key steps of the preprocessing workflow, starting from barcodes identification up to the generation of gene and isoform count matrices. First, we compared their running time and memory usage, and tested their suitability for large datasets. Then, we compared the number of UMIs and genes detected with paired short-read data. Subsequently, we evaluated the ability of methods to identify the cell-associated barcodes, extract and correct UMI tags. Next, we tested the fidelity of gene expression quantification at both single-cell and pseudo-bulk levels using simulated data. Then, we assessed the quality of cell type annotation obtained from real long-read data, as well as similarity and integration with short-read data. In addition, we evaluated the quality of spatial clustering by assessing the spatial continuity of the clusters and the similarity with clustering obtained from short-read data. Finally, we evaluated the ability of methods to correctly assign and quantify known isoforms, to discover novel ones, and to detect differential isoforms usage between cell clusters. The main results are summarized in Figure 2.

**Fig. 2.**
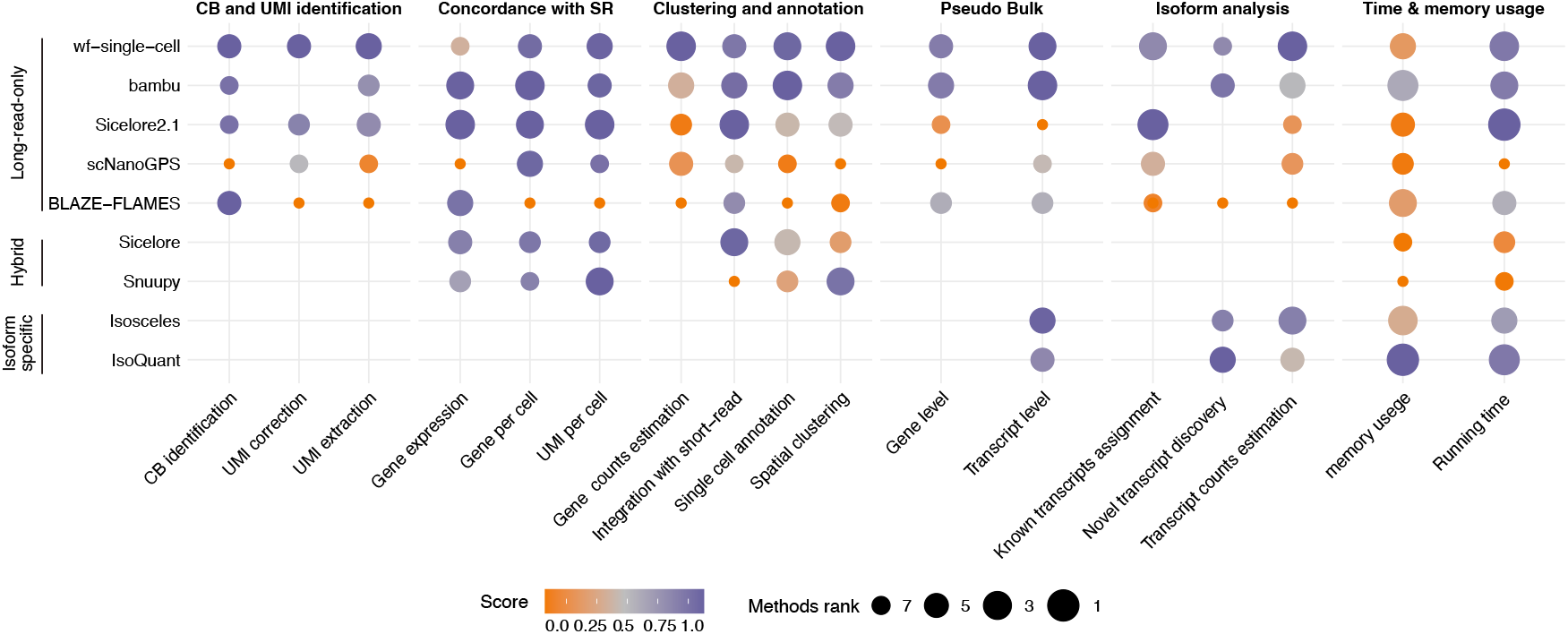
Ranking of methods across key aspects of evaluation —. Methods are ranked by overall score in each category (long-read-only, hybrid and isoform-specific). Metrics are divided into barcode and UMI identification, concordance with short-read data, and biological results at single-cell and bulk levels and isoform analysis criterion. Methods are scored by key analytics metrics. Each metric was scaled to the interval [0,1] and harmonic means are computed across datasets to obtain a score. Overall benchmarking scores are the computed mean of all scores (see Methods). The colour indicates the score and size of the circle indicates ranking of methods, where a large blue circle represents the best rank. Missing dot indicates whether an aspect is not covered by the method or the measurement was not able to be obtained.

Our benchmarking workflow is provided as a reproducible Nextflow pipeline named scKeñver. It allows developers to test and evaluate new methods using our or their own datasets. All analyses were implemented in executable and traceable R notebooks. The computational reproducibility is enabled by using the Docker containerization technology (44).

### Running time and memory usage

We compared the runtime and memory usage of the ten evaluated methods (Figure S1), which showed differences between hybrid, long-read-only and isoform-specific approaches.

While the two hybrid methods, Snuupy and Sicelore, required respectively 25 and 11 hours, to process 10 millions reads, almost all long-read-only methods took less than 4 hours (Sicelore 2.1: 1h25min, Bambu: 1h39min and wf-single-cell: 1h45min, BLAZE-FLAMES: 3h30min). Exception is made by scNanoGPS, which exhibited a remarkably longer runtime than other long-read-only methods (20h25min), primarily due to its barcode curation step (16h58min with 30 threads) (Figure S1a). Moreover, the curation step of scNanoGPS is heavily influenced by the size of the barcode library: 9h23min for 900 cell-associated barcodes vs 34h41min for 4,500 cell-associated barcodes for a total of 1 million reads in both datasets (Figure S1b). Regarding memory usage, the three long-read-only methods FLAMES, wf-single-cell and Bambu demonstrated remarkable memory efficiency compared to hybrid methods. However, the amount of memory used by Sicelore 2.1 and scNanoGPS was comparable to hybrid methods (Figure S1c). Concerning isoform-specific methods, IsoQuant outperformed Isosceles both in terms of running times (15 min versus 57 min to process 10 millions reads) and memory usage (26 MB versus 8GB for 10 millions reads).

Considering scalability, Snuupy did not succeed in processing the PromethION datasets, as it required up to 200 GB of RAM for BLAST step. This poor scalability makes it unsuitable for large-scale datasets, without improvements. Similarly, scNapBar failed to produce results with MinION datasets (about 10 million reads) even after 120 hours of running. It was thus not tested on other datasets and was excluded from subsequent evaluations.

### Comparison of the gene count matrices obtained from long-read and short-read scRNA-seq data

To evaluate the gene count matrices obtained after preprocessing the long-read datasets, we first assessed the number of UMIs and corresponding genes identified per cell or spot across methods. The raw gene expression matrix output of each method was used to quantify (i) the number of UMIs per barcode, (ii) the number of genes detected per barcode (genes with non-zero expression), and (iii) the total number of UMI counts in the matrix. These metrics were then compared to short-read data. Furthermore, we compared the gene expression for each barcode between short- and long-read data (Figure 3).

**Fig. 3.**
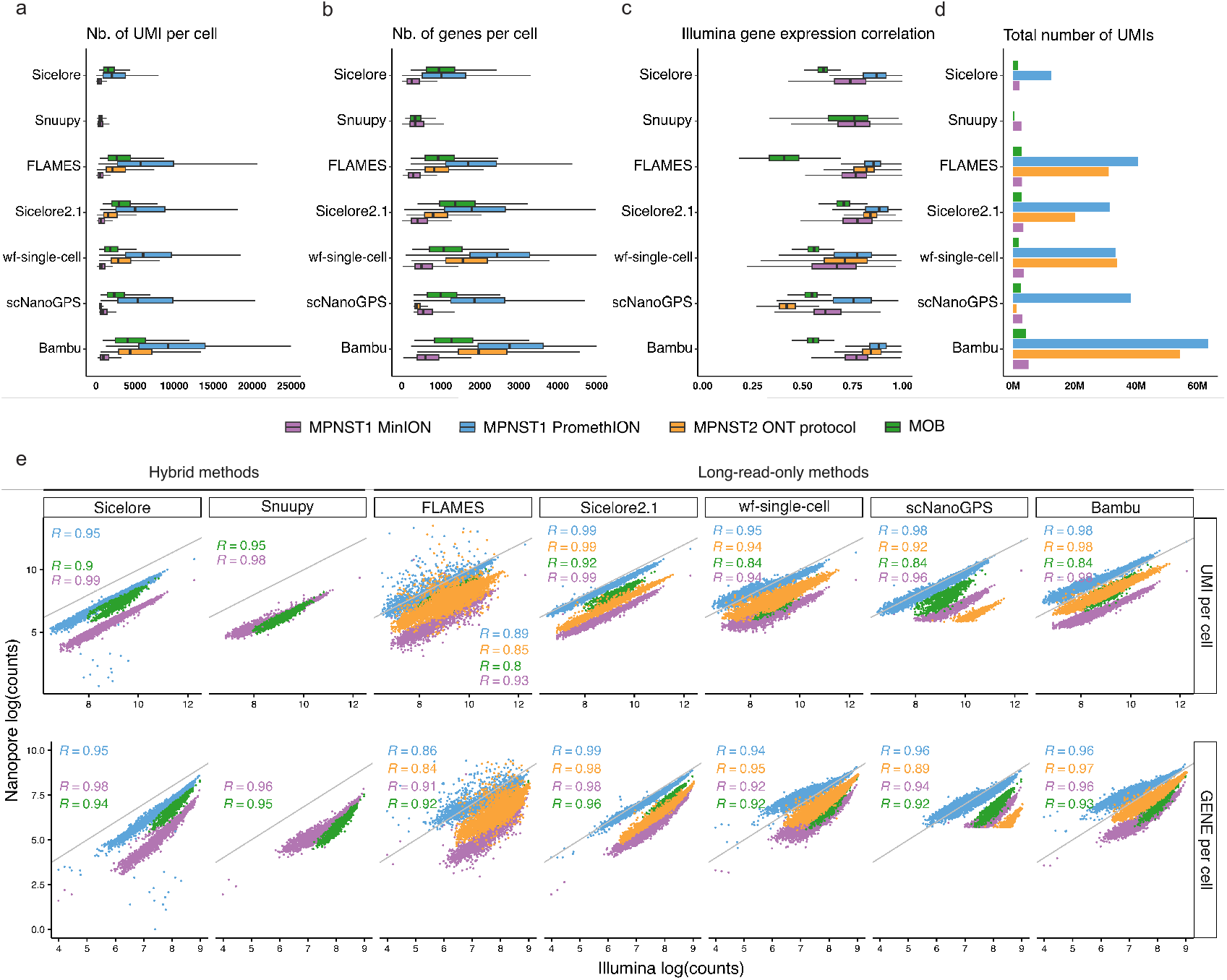
Comparison of gene count matrices obtained from long-read versus short-read data —. The gene count matrices were obtained by processing the four Nanopore long-read datasets (see legend with associated colors) with each bioinformatics method listed in the y axis (panels a-d). Isosceles and IsoQuant are not included as they do not process reads into count matrices. **(a)** Number of UMIs per barcode. **(b)** Number of genes per barcode. **(c)** Box plot showing the distribution of Pearson correlation coefficient (r) of the number of UMI per gene between short and long read data. r values are calculated for each barcode, boxes represent the 25% quantile to 75% quantile range and the median. **(d)** Total number of detected UMIs per method. For runtime reasons, the MPNST2 dataset prepared with the sc-ONT standard protocol (in orange) was solely processed with the five long-read-only methods. **(e)** Scatter plots showing the number of UMI and number of genes per barcode (log-normalized values) detected in short read (x-axis) and long read (y-axis) data. The Pearson correlation coefficient for each dataset is shown in the same color. Gray lines represent the x=y relationship.

Distinct trends emerged across the computational tools, particularly in the total number of UMIs detected, and consequently in the number of UMIs per barcode. Long-read-only approaches tend to detect a greater number of UMIs compared to hybrid approaches (Figure 3a-d), which may reflect both increased sensitivity and/or a higher rate of false positives. To distinguish between these two cases, we used the short-read dataset as a reference. With respect to UMI and gene detection per cell, Bambu showed a slight increase in UMI counts while maintaining high correlation with short-read data, suggesting greater sensitivity in UMI detection (Figure 3a-c). The elevated number of detected genes in Bambu can also be attributed to its ability to predict novel genes, contributing to its expanded gene detection profile.

We then compared gene expression with short-read data using Pearson’s correlation. Overall, we found relatively high average Pearson correlations with all tools (Figure 3c). We observed consistent trends across data types and sequencing depth. To evaluate the effect of sequencing depth, we compared the same sample sequenced on high-versus low-throughput platforms. Notably, all workflows performed better with higher sequencing depth (MPNST1 PromethION data) compared to lower sequencing depth (MPNST1 MinION data), attesting a depth sensitivity. In contrast to the scRNA-seq datasets, which exhibited high median Pearson’s correlation coefficients, the spatial transcriptomics data showed substantially lower correlations. Specifically, Pearson’s correlation coefficients ranged from r=0.4 for FLAMES to r=0.75 for Snuupy, indicating poor concordance. These values are even lower than those observed with the MinION scRNA-seq data, despite its lower sequencing depth and saturation. These results suggest that on this dataset, either low sequencing depth and/or spot resolution of the Visium platform may compromise the accurate representation of the transcriptome.

Finally, we compared the number of UMI and genes detected within each cell, between long-read and short-read data (Figure 3e). The same conclusions are reached, either considering the number of UMI or the number of genes. High Pearson correlations were observed for all methods, highlighting that long-read and short-read data are consistent to evaluate the biology of cells. By comparing computational methods, lower correlations are found using FLAMES. This can be explained by the UMI correction issues, associated with sequencing errors. By comparing biological protocols, lower correlations are found when evaluating the number of UMI per spot, in spatial data, using the long-read-only methods (Figure 3e, green dots). Coherently, regardless of the protocol, correlations are higher when considering hybrid methods than long-read-only methods. This is expected, as, by definition, hybrid methods rely on paired short-read data for UMI assignment. However, although correlations are high, hybrid methods tend to detect less UMI and genes per cell. Similarly, when comparing sequencing platforms, we observed underestimation of both the number UMI and the number of genes per cell using MinION, compared to PromethION. This can be related to the lower sequencing depth associated with MinION flowcells.

### Accuracy in barcodes/UMI assignment and gene quantification fidelity

We evaluated the performance of the five long-read-only tools—BLAZE-FLAMES, Sicelore 2.1, wf-single-cell, scNanoGPS and Bambu—in accurately identifying barcodes and correcting UMI sequences, as well as their fidelity and accuracy in quantifying gene expression at both the single-cell and pseudo-bulk levels. The barcodes identification consists in intersecting the barcodes inclusion list (formerly called whitelist), provided by 10X Genomics, with the set of the raw barcodes obtained from the sequencing. This intersection is called the shortlist and corresponds to potential cells/spots. From this shortlist, cell-associated barcodes are identified from the kneeplot.

To assess their ability to detect cell-associated barcodes, we first compared the barcode list produced by each tool to the set of barcodes identified from the paired short-read data on the MPNST1 PromethION dataset (Figure 4a). In total, we identified 4,513 cell-associated barcodes in the short-read MPNST1 dataset. Among these, 3,903 barcodes (92%) were recovered by all long-read-only methods (Figure 4a). Notably, scNanoGPS, Sicelore 2.1, and Bambu reported additional cell-associated barcodes not present in the short-read shortlist, respectively 1,465, 484, and 220 extra barcodes. Comparable performance patterns were observed in both the ScNaUmi-seq MinION and ONT PromethION datasets (Figure S2a-b).

**Fig. 4.**
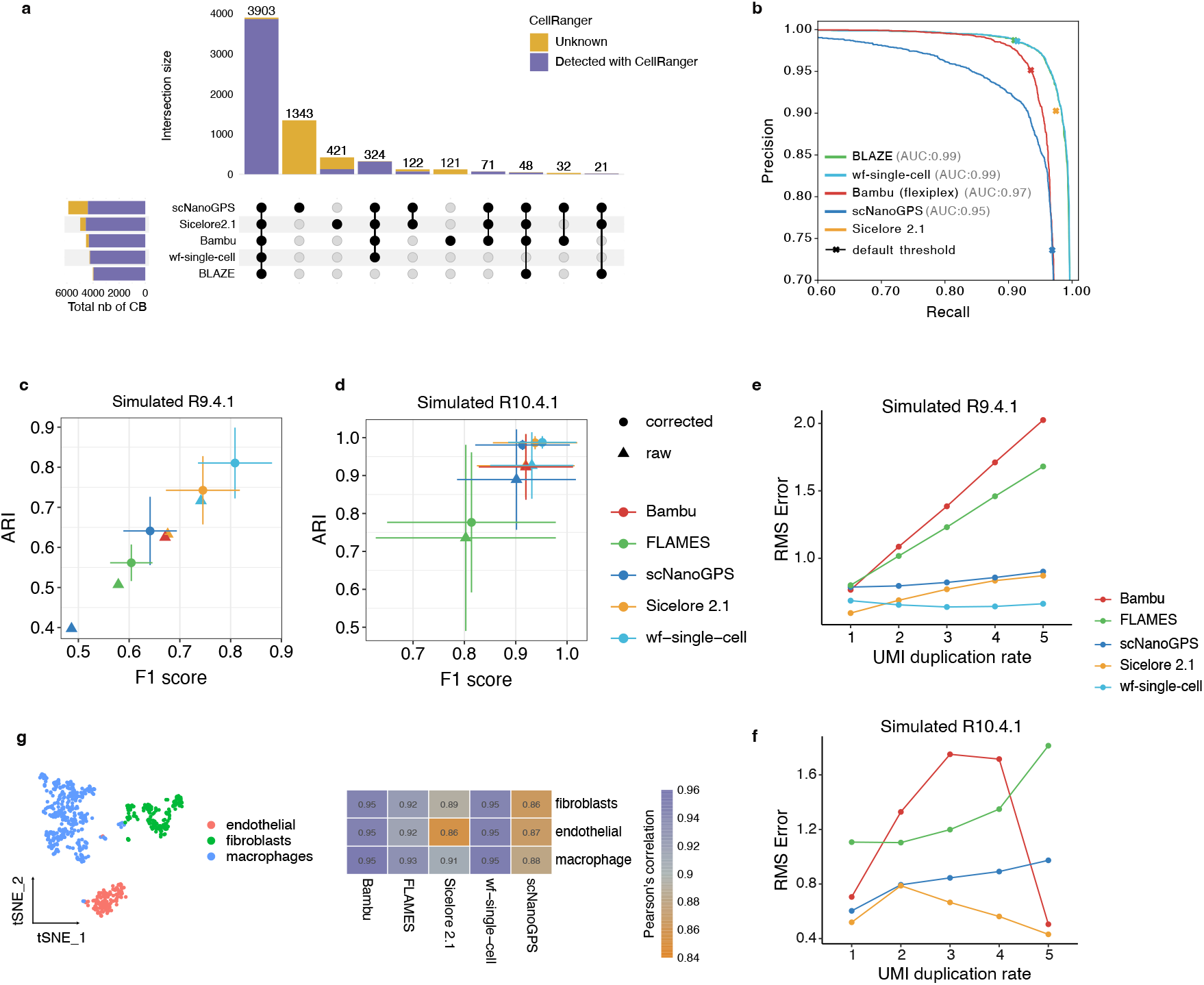
Barcode, UMI, and transcript identification —. **(a)** Barcode upset plot comparing different cell-associated barcode lists. The bar chart on the left shows the total number of barcodes found by each tool. The bar chart on top shows the number of barcodes in the intersection of barcode lists from specific combinations of methods. The dots and lines underneath show the combinations. **(b)** Precision-recall curves across different count thresholds. Precision and recall were calculated across different count thresholds by defining the barcodes identified from short reads as the ground truth, specifically the shortlist from Cell Ranger after the removal of empty droplets. **(c-d)** Scatter plots of the mean F1-score against mean ARI score for the UMI error correction. Error bars indicate the mean and standard error across simulated R9.4.1 **(c)** or R10.4.1 **(d)** datasets, on which methods run (n=2; UMI duplication 1 and 5). **(e-f)** Comparison of gene expression estimation by different methods on the simulated R9.4.1 **(e)** or R10.4.1 **(f)** datasets (1 million reads each) across five different UMI duplication rates, evaluated by root of mean square error. **(g)** (left) UMAP projection of gene expression level quantifications from ground truth of simulated data, colored by cell type. (right) Pearson’s correlation (color scale) of each method’s quantifications with expected quantification at pseudo-bulk level.

Since BLAZE, wf-single-cell, scNanoGPS, and Bambu infer the set of cell-associated barcodes by identifying an inflection point in the ranked barcodes count distribution, we computed precision–recall curves across varying barcode counts threshold to evaluate the false-positive detection rates of these approaches. As a ground truth reference, we used the barcode list generated from the short-read dataset by CellRanger, after filtering out empty droplets using the DropletUtils R package (45) (Figure 4b). BLAZE and wf-single-cell showed the best performances (precision=0.91; recall=0.99 on default threshold) with a high consistency in the detection of cell barcodes, minimizing the false-positive. They are followed by Bambu (precision=0.95; recall=0.94 on default threshold). Although Sicelore 2.1 uses a different strategy to define the barcode shortlist, we compared its performance with the other methods. Unlike other long-read-only methods, Sicelore 2.1 does not rely on the inflection point of the ranked barcodes count distribution. Instead, it merges the sequenced barcodes with all possible 10X Genomics barcodes (inclusion list) within a defined Levenshtein distance. This strategy prioritizes a high recall (0.97) but leads to a slightly higher false-positive rate (precision=0.9) (Figure 4b). The lower performances were observed with scNanoGPS (precision=0.74 and recall=0.97 on default threshold) that assigns barcodes without any prior knowledge of the 10X Genomics barcode inclusion list (Figure 4b).

Next, we evaluated the performance of the UMI error correction strategies of each method. To obtain a reliable ground truth, we simulated single-cell long-read datasets using the AsaruSim tool (42). Five datasets, each consisting of *~*1 million reads, were generated with a median UMI duplication rate ranging from 1 to 5 duplications per UMI and achieving a sequencing saturation of 20%, 35%, 45%, 54%, and 60%, respectively (Table S1, R9.4.1 dup1-dup5). To generate realistic reads, our real datasets (MPNST1 PromethION with R9.4.1 Nanopore chemistry) were used as a reference model to replicate the error profile (median phred score 13). Additionally, to investigate the impact of sequencing quality in the performances of tools, we simulated 5 other datasets using a sequencing error model trained on the latest R10.4.1 Nanopore chemistry (median phred score 20) (Table S1, R10.4.1 dup1-dup5). After preprocessing the simulated datasets, we obtain raw UMI sequences and corrected UMI sequences resulting from a UMI error correction step. The corrected UMI are compared to the true UMI sequences (ground truth set in the simulation), measured with an F1-score. To assess the ability of each method to cluster UMIs originating from the same original transcript, we calculated the Adjusted Rand Index score (ARI) (Figure 4c-d). Considering the R9.4.1 Nanopore chemistry, wf-single-cell exhibited the best UMI clustering performance, achieving a mean F1-score of 0.8 and the highest ARI score of 0.81. Sicelore 2.1 followed, with a mean F1-score of 0.74 and an ARI score of 0.74, while Bambu and FLAMES showed the worst performance F1-score 0.67 and 0.6 and ARI score 0.62 and 0.56 respectively (Figure 4c, circle-shaped dots). Similar trends were observed with the R10.4.1 datasets, with an overall increased performance of the tools reaching F1-score of 0.95 and an ARI score of 0.98 (Figure 4d).

To further assess eventual variations in the efficiency of UMI sequence searching strategies, we compared the raw UMI sequence with the true UMI sequences and computed the same metrics (Figure 4c-d, triangle-shaped dots). For both R9.4.1 and R10.4.1 datasets, we observed significant differences among tools in terms of the quality of raw UMI sequences. wf-single-cell demonstrated the highest quality of raw UMI sequences (F1-score=0.74; ARI=0.71), whereas scNanoGPS exhibited the worst UMI searching performance in R9.4.1 data (F1-score=0.48; ARI=0.39). The large difference in the quality of scNanoGPS raw and corrected UMI, especially in high error conditions (R9.4.1 datasets), suggests inferior UMI searching but good correction strategy. These findings suggest that the initial UMI sequence search and identification also plays a crucial role in achieving effective UMI deduplication.

Next, we investigated the impact of UMI deduplication on gene expression quantification. To this end, we compared the gene expression matrix generated by each tool to the expected (ground truth simulation) gene expression counts. To ensure consistent comparisons across tools and to reduce biases arising from library size, we applied a normalization step in which each matrix was standardized using the mean and standard deviation derived from the expected count matrix. Then, the fidelity of gene quantification across UMI duplication, measured as sequencing saturation, was assessed by computing the mean squared error (MSE) between estimated and expected gene expression values (Figure 4e-f). Sequencing saturation reflects the fraction of reads that are UMI duplicates, i.e., originated from already-observed UMI. We found that increasing sequencing saturation leads to a gradual increase in gene expression estimation error for some methods (Figure 4e). Notably, wf-single-cell showed the lowest estimation error across duplication rates (MSE=0.66 at 5 UMI duplications), while FLAMES and Bambu exhibited substantial increases in MSE (exact values: MSE=1.6 and 2.02, respectively, at 5 UMI duplications). Similar trends were observed in the R10.4.1 dataset, where wf-single-cell achieved the best estimation accuracy (MSE=0.43 at 5 UMI duplications), followed by Bambu (MSE=0.5 at 5 UMI duplications) (Figure 4f). Of note, Bambu performs UMI deduplication without a UMI error correction step. Cumulatively, its UMI deduplication relies on a fixed threshold for the number of UMIs, which makes its performance highly dependent on sequencing depth and read quality. This explains the sharp increase of performance at a 5 UMI duplication rate in the R10.4.1 simulated data (Figure 4f).

Overall, these results highlight the strong correlation between effective UMI error correction and accurate gene expression quantification. Indeed, tools with robust UMI correction—such as wf-single-cell and Sicelore—showed the most faithful gene-level estimates at high sequencing error rates. To support this observation, we simulated 20 million reads from 630 cells across three cell types—macrophages, endothelial cells, and fibroblasts—derived from the MPNST1 datasets, using two ONT error models (Table S1, Simulated R9.4.1 and Simulated R10.4.1 datasets) (Methods). Overall, wf-single-cell yielded the best gene count estimations in both chemistries (MSE=0.44 in R9.4.1 and MSE=0.32 in R10.4.1), followed by Sicelore 2.1 (MSE=0.63 in R9.4.1) and Bambu (MSE=0.58 in R10.4.1). These results confirm Bambu’s marked performance gain specifically under the 5-UMI duplication condition in the R10.4.1 dataset (Figure S2c-d).

Next, we investigated the gene expression at the pseudo-bulk level. We compared the estimated gene expression counts from each cell type with the true gene expression counts (Methods). Bambu and wf-single-cell achieved the best Pearson’s correlation for the three cell types (r=0.95), followed by FLAMES (r=0.93, 0.92, 0.92 for fibroblast, endothelial cells and macrophages respectively) and Sicelore 2.1, achieving the lowest correlation (r=0.91, 0.86, 0.89) (Figure 4g). Similar results were observed with the R10.4.1 chemistry (Figure S2e).

### Evaluation of biological results at the gene level

To benchmark the effect of processing workflows on the downstream analysis results at the gene level, we first focused on the cell projection and cell type annotation with the MPNST1 PromethION dataset, on which long-read methods showed the best performances. As reference, the short-read sequencing of this dataset captured about 4,500 cells, corresponding to 14 cell types dominated by a large population of tumor cells. All gene count matrices preprocessed by CellRanger (short reads) or the tested tools (long reads) are analysed similarly: after count matrix processing using Seurat V5, cells passing quality control steps were retained, annotated for cell types, and projected using UMAP (methods). Regardless of the methods, the MPNST1 PromethION dataset contains four main populations, annotated as tumor cells (main population), macrophages, fibroblasts and endothelial cells (Figure 5a).

**Fig. 5.**
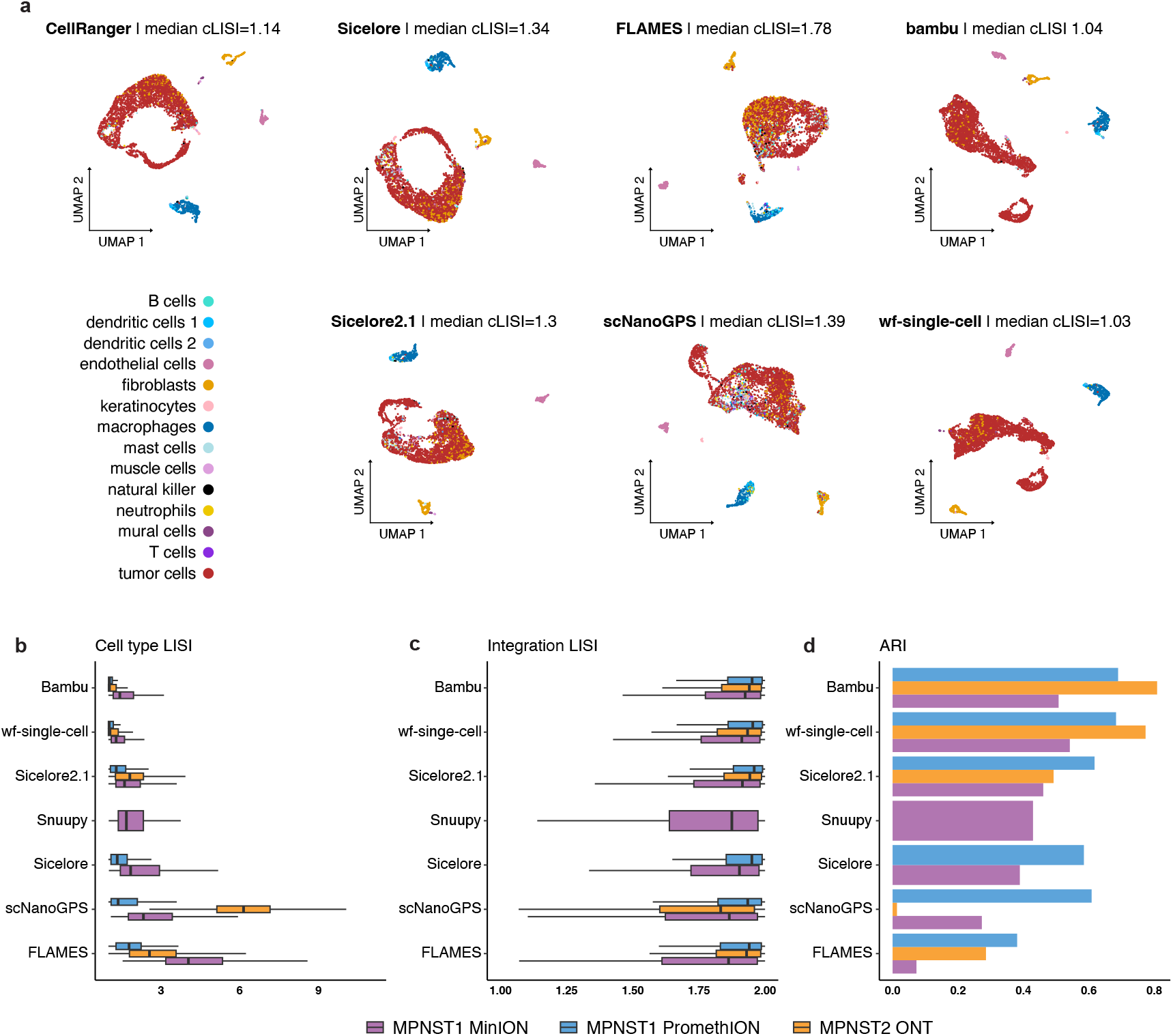
Biological results and comparison with short read data at the gene level —. **(a)** UMAP representation of the MPNST1 PromethION dataset with cells colored by cell type annotation. The CellRanger UMAP visualization is based on short-read data, all other methods are based on long-read data. The median cell type LISI (cLISI) is indicated at the top of each representation. **(b)** Quantitative assessment of cell type annotation performances in each method as measured by cLISI. Lower values indicate homogeneous annotation. **(c)** Quantitative assessment of alignment of UMAP representation of long-read versus short-read datasets as measured by integration LISI (iLISI). **(d)** Quantitative assessment of the concordance of long-read based with short-read based annotation cell type annotation measured by ARI score. Datasets used in b-d are color coded according to the legend below the panels.

To assess the quality of the cell-type annotation between short-read data processed with CellRanger and long-read data processed using various computational methods, we used an independent metric. Cell-type local inverse Simpson’s index (cLISI) quantifies the degree of local cell-type separation by combining the cell type annotation and the cell coordinates, with lower values indicating more homogeneous and consistent labeling (46). wf-single-cell provided the most accurate cell-type characterization, both visually (Figure 5a) and quantitatively, with a mean cLISI of 1.03 (Figure 5b). It was followed closely by Bambu (1.04), then Sicelore 2.1 (1.30), Sicelore (1.37), scNanoGPS (1.39), and FLAMES (1.78), as measured on the MPNST1 PromethION dataset. Notably, both wf-single-cell and Bambu outperformed the short-read reference (median cLISI < 1.14), suggesting that long-read gene expression profiling can achieve more precise cell-type separation than traditional short-read approaches. Consistent results were observed across MPNST1 MinION and MPNST2 ONT datasets (Figure 5b).

To further compare long-read data processing by considering short-read data as reference, we merged each pair of long-read and short-read count matrices and generated a batch-effect corrected projection (methods). Then, we computed two complementary metrics. The integration local inverse Simpson’s index (iLISI) measures the degree of dataset mixing in a shared embedding space (46). Secondly, the Adjusted Rand Index (ARI) assesses the consistency of cell-type annotations between the long-read and the short-read data. Bambu achieved the highest median iLISI and ARI scores across datasets (median iLISI=1.94; ARI=1.66), indicating better alignment and agreement with the short-read reference (Figure 5c-d), followed by wf-single-cell (median iLISI=1.93; ARI=1.66) and Sicelore 2.1 (median iLISI=1.94; ARI=1.52). In contrast, FLAMES (median iLISI=1.92; ARI=1.24), and scNanoGPS (median iLISI=1.90; ARI=0.29) showed weaker agreement, with slightly lower integration and clustering consistency.

We further investigated some genes with expression detected in long-read but not short-read datasets. We observed that incomplete 3’-UTR annotations hampers the quantification of the 3’ signal in short-read data, resulting in missing or misquantified gene expression values (Figure S3). These biases can distort transcriptomic profiles and result in the misclassification of cell types. In contrast, as long-read sequencing is not restricted to 3’ end, it enabled accurate gene expression quantification despite incomplete 3’-UTR annotations, enhancing the cell type identification.

Regarding spatial transcriptomics, the MOB Illumina dataset from the mouse olfactory bulb (Figure 6a), processed with the 10X Genomix Space Ranger pipeline, resulted in 916 spatial spots detected. Unsupervised clustering identified six clusters, consistent with the known anatomical regions of the olfactory bulb (41). To assess gene-level performance in long-read methods, we preprocessed the MOB PromethION dataset with each method, and then performed similar unsupervised clustering of the detected spots. When comparing the clustering results to the short-read reference using ARI, wf-single-cell showed the highest similarity (ARI=0.75), followed by Bambu (ARI=0.63) and Sicelore (ARI=0.59) (Figure 6b). We further evaluated spatial clustering continuity using cLISI and CHAOS scores (47, 48), where lower values reflect more spatially coherent clusters. wf-single-cell again produced the most continuous and homogeneous spatial clusters (median cLISI=1.01; CHAOS=0.12), followed by Sicelore (cLISI=1.05; CHAOS=0.12) (Figure 6c-d).

**Fig. 6.**
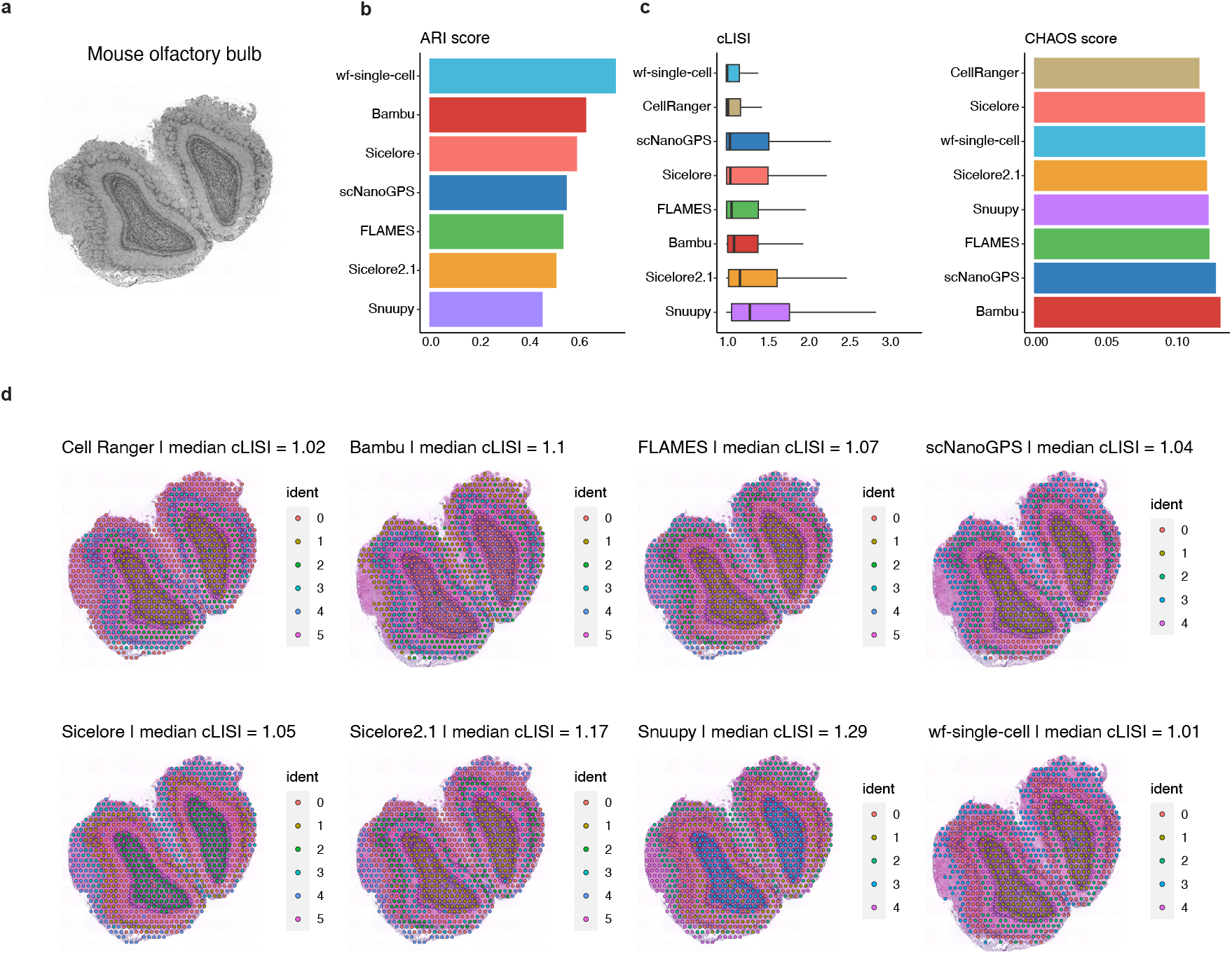
Comparison of spatial data clustering with short read data at the gene level —. **(a)** Slice of the mouse olfactory bulb, obtained by SpaceRanger from the repository (Accession number GSE153859). **(b)** Quantitative assessment of concordance of spatial clustering obtained with long-read (MOB PromethION) versus short-read (MOB Illumina) datasets, as measured by ARI score. **(c)** Quantitative measurement of the quality of spatial clustering using cLISI and CHAOS metrics, the lower the value, the more the method yields a continuous and homogeneous neighborhood for spatial spots. **(d)** Comparative visualization of clusters identified by long-read methods.

Together, these results demonstrate that long-read sequencing can achieve high-resolution clustering and accurate cell-type annotation, both in scRNA-seq and spatial transcriptomics data. Among the tested methods, wf-single-cell, Bambu followed by Sicelore consistently provided the most biologically meaningful clustering patterns, and strong performance in preserving biological signals during preprocessing.

### Evaluation of biological results at the isoform level

#### Isoforms characterization and quantification

To benchmark transcript assignment accuracy, we evaluated the ability of each long-read-only method to correctly assign simulated long reads to annotated transcript models. To this end, we compared the annotated transcript model assigned to each read with its corresponding ground-truth transcript label in the Simulated R.9.4.1 dataset (Figure 7a). Since Isosceles, IsoQuant and Bambu assign transcript only after the UMI deduplication step, it was not possible to directly evaluate transcript assignment at the individual read level for these two methods. Sicelore 2.1 achieved the highest performance in transcript assignment (F1-score=0.88), likely due to its ‘strict’ mode, which requires a complete exon-exon structure match for assignment. This was followed by wf-single-cell (F1-score=0.78), while scNanoGPS and FLAMES showed more modest performance, with F1-scores of 0.54 and 0.39, respectively.

**Fig. 7.**
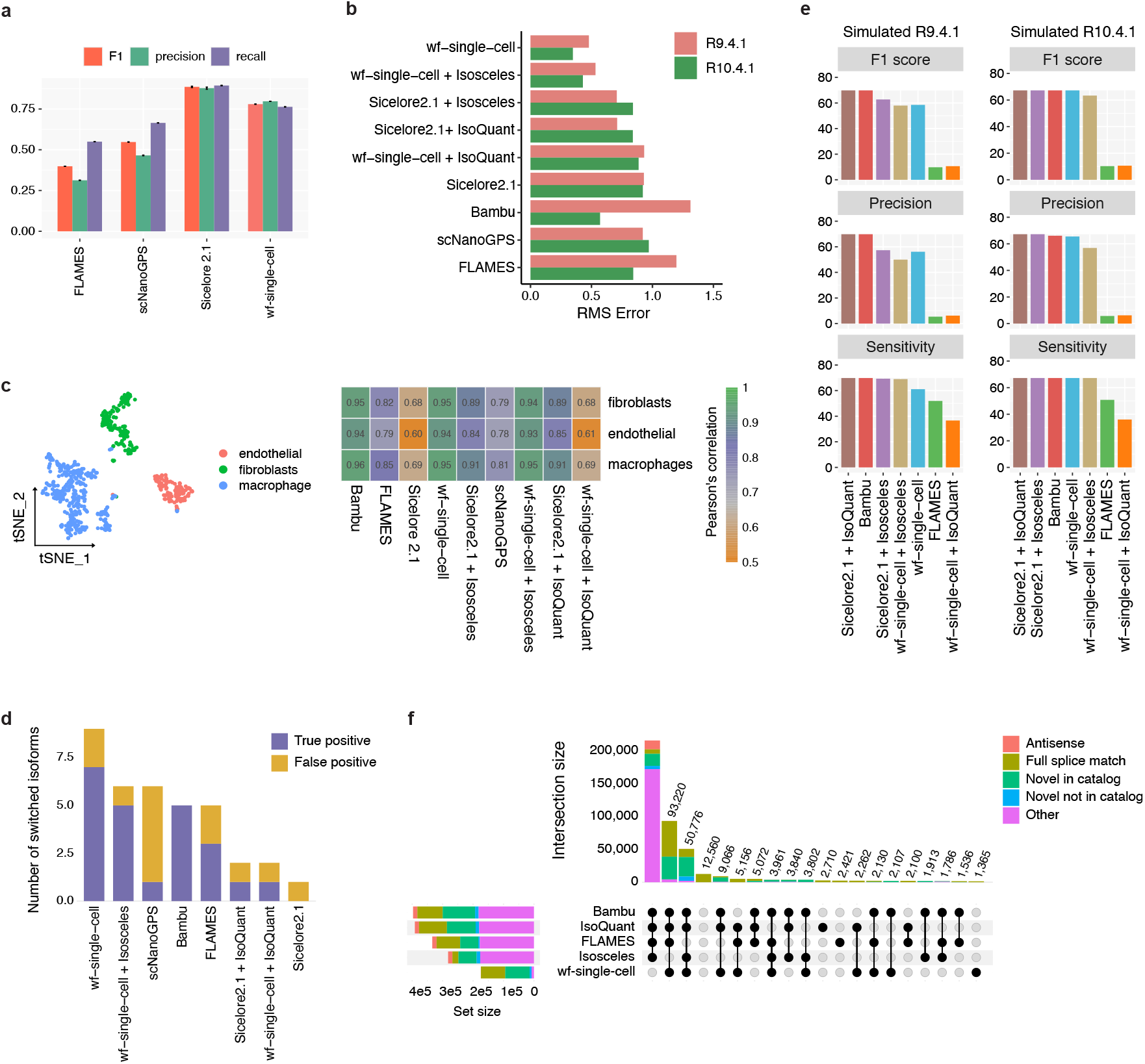
Isoforms characterization, quantification and discovery —. **(a)** Ability of the tested tools to assign reads to annotated transcripts, as measured using precision, recall and F1-scores on the Simulated R9.4.1 dataset. Metrics are computed by comparing each transcript label assigned to reads with the labels of the ground truth. **(b)** Ability to quantify transcripts measured using RMSE between the observed and expected (ground truth) transcript expression values. RMSE were computed after a normalization step, where each isoform count matrix was standardized using the mean and standard deviation of the ground truth. Both Simulated R9.4.1 and Simulated R10.4.1 datasets are shown. **(c)** (left) tSNE projection of the Simulated R9.4.1 dataset at the isoform level. Cells are colored by cell type annotation. (right) Pearson’s correlation coefficient (color scale) of each method’s quantifications with the expected qualifications across three cell types, while considering pseudo-bulk. **(d)** Barplot representing the number of differentially expressed isoforms between the three clusters identified by each tool, and whether they are present in the expected list (true positive) or not (false positive). Note that Sicelore 2.1 + Isosceles did not detect any significant switched isoforms and is not represented on the barplot. **(e)** Precision, sensitivity and F1-score rates of novel isoforms discovery by each tool in R9.4.1 and R10.4.1 simulated data. **(f)** Upset plot showing the 20 largest intersections of isoform coordinates between isoform annotations identified by each tool in the real MPNST1 PromethION data. The bars are colored by isoform structural categories.

Next, we assessed the performance of these tools in transcript-level quantification by comparing the estimated transcript abundances with the expected values in the Simulated R9.4.1 and Simulated R10.4.1 datasets. As with gene expression analysis, we computed the root mean squared error (RMSE) after a normalization step, where each isoform count matrix was standardized using the mean and standard deviation of the ground truth (Figure 7b). In both simulated datasets, wf-single-cell, either alone or coupled with Isosceles, achieved the most accurate isoform quantification, with the lowest RMSE values (0.48 and 0.53 in R9.4.1; 0.35 and 0.43 in R10.4.1, respectively). These were followed by Sicelore 2.1 coupled with Isosceles (RMSE=0.71 in R9.4.1 and 0.84 in R10.4.1), while scNanoGPS and FLAMES exhibited the highest deviations, with RMSE values of 0.92 and 1.2 in R9.4.1, and 0.97 and 0.84 in R10.4.1, respectively. These results consistently demonstrate the high fidelity of wf-single-cell in quantifying known transcript isoforms across both sequencing chemistries.

Transcript quantification was further evaluated at the pseudo-bulk level by aggregating isoform counts across cells within the three simulated cell types. Overall, we observed a high correlation between the resulting expected and observed transcript abundances (Figure 7c), supporting the ability of long-read methods to reliably quantify isoforms at the population level. Bambu exhibited the highest concordance with the ground truth, with Pearson correlation coefficients of 0.95, 0.94, and 0.96 for fibroblast, endothelial cells, and macrophage populations, respectively. This was followed by wf-single-cell and Isosceles, which also showed strong correlations. In contrast, Sicelore 2.1 demonstrated the lowest correlation scores among the evaluated methods. Similar trends were observed in the R10.4.1 dataset (Figure S2f), further confirming the consistency of these findings across different sequencing chemistries.

#### Differential isoform analysis

Next, we investigated genes affected by differential isoform usage between cell types with the Isoswitch R package that leverages the Seurat’s FindMarkers function to identify isoform switches between each possible pair among the three cell populations of the simulated datasets. Differentially splice genes (DSGs) were filtered using an adjusted p-value threshold of 5%. We then compared the list of DSGs from long-read methods to the ground truth (Figure 7d). Bambu led in performance, identifying 10 out of 18 DSGs confirmed by the ground truth. wf-single-cell combined with Isosceles followed closely with 8 out of 17 confirmed isoforms, and FLAMES identified 9 out of 14. In contrast, Sicelore 2.1 alone did not detect any confirmed DSGs.

We then analysed the real MPNST1 PromethION dataset to evaluate the effect of each preprocessing tool on the final detection of DSGs between cell clusters. This analysis was performed on each isoform count matrix with Isoswitch, between each pair of the 10 cell clusters, to identify the genes showing an isoform switch between two cell clusters (Figure S4a). Few isoforms are detected consistently by four tools (green bars). Strikingly, Isosceles, FLAMES and scNanoGPS lead to the isolated detection of 2,124, 1,580 and 624 switched isoforms respectively, potentially false positives. This large number of potential false detection can be due to transcript assignment or quantification errors. To investigate the biological functions behind the detected isoform switch, we performed gene ontologies analysis using the 2,942 genes detected by all methods (Figure S4b). Genes characterized by isoforms switches are mostly related to the processing of RNA molecules, suggesting that they are themselves directly related to the splicing process. Among the top ten genes characterized by isoforms switches, consistently detected by at least six tools, we pinpointed *Cdkn2c*, as in this biological context, *Cdkn2a* is known to be deleted (49) (Figure S4c). Additionally, other isoforms switches concerned variations between cell types, the isoform switch of *Cdkn2c* was restricted to the tumor cell population (Figure S4d-e, in purple). We observed that the expression of the isoform 202 coincides with a proliferative cell activity (Figure S4f), while the isoform 201 is over-expressed in quiescent cells. This observation is consistent with observations made in T cells in physiological conditions (50). Very recently, alternative splicing of *Cdkn2c* has been shown to be correlated with distinct response to treatment in the context of ovarian cancer (51).

#### Isoforms discovery

Among the evaluated tools, only wf-single-cell, FLAMES, Isosceles, and Bambu support transcript discovery, i.e. novel transcripts not part of the annotation (Table 2). We compared their performances on the Simulated R9.4.1 and Simulated R10.4.1. Transcript discovery was evaluated by randomly removing 4,412 (15%) expressed isoforms-defined as isoforms with non-zero UMI counts-from the GENCODE annotation, to mimic real-world scenarios where unannotated transcripts are present (Methods). This reduced annotation was then provided as the reference for each tool, and their ability to recover the removed transcripts as novel was assessed using gffcompare (52) (Figure 7e). The set of predicted novel transcripts from each tool was compared to the set of intentionally removed isoforms to evaluate transcript discovery performance.

Compared to most tools, Bambu alone yields highly accurate novel transcript models on R9.4.1 data, likely due to its splice alignment correction strategy, achieving an F1-score of 73%. Isosceles demonstrates slightly higher precision (57%) when coupled with Sicelore 2.1 compared to its combination with wf-single-cell (50%), while maintaining a comparable sensitivity (69%) in both cases. Isosceles combined with Sicelore 2.1 reports 1,083 (15%) false positive exons, compared to 2,361 (14%) when used with wf-single-cell. This difference is likely attributable to splice site alignment errors, suggesting improved accuracy when using the consensus-based correction methods employed by Sicelore 2.1. On the R10.4.1 dataset, Isosceles slightly outperformed other methods, achieving an F1-score of 70%, compared to 68.5% for Bambu, 68% for wf-single-cell, and 63% for Isosceles coupled with wf-single-cell. In contrast, FLAMES showed poor recall across both datasets, highlighting its limitation in novel isoform detection.

To examine transcript assembly on a real dataset, where the ground truth is unknown, we compared the coordinates of all detected isoforms with the known transcripts from the reference annotation. On the MPNST1 PromethION dataset, the highest percentage of known transcripts, represented by the ‘full splice match’ isoform structural category detected by gffcompare, was achieved by wf-single-cell (79,986 transcripts; 46%) (Figure 7f). The number of novel transcripts, represented by all ‘novel’, and ‘other’ categories, shows the highest differences. While wf-single-cell had the lowest number of ‘other’ isoform structural categories (4%), a very high percentage of novel transcripts were detected by Bambu (262,175 transcripts; 93%) and IsoQuant (256,260 transcripts; 77%) (Figure 7f). Regarding the intersection between annotations, Bambu and IsoQuant achieve the highest number of isoforms confirmed by at least three other tools (363,755 and 359,810 isoforms), wf-single-cell achieves the lowest number (147,973 isoforms). Similar trends are observed regarding only isoforms reported as novel (Figure 7f). These results are consistent with our observations from the simulated data, suggesting the high reliability of Bambu and IsoQuant as isoform discovery methods.

## Discussion

We benchmarked 10 single-cell and spatial long-read tools using several datasets generated with different sequencing platforms, protocols and chemistries. We used several metrics that measure quality control and concordance with short-read data, including the accuracy in barcodes/UMI assignment, the fidelity of gene and transcript expression quantification, the clustering and cell type identification at single-cell and spatial level as well as the isoforms characterization and quantification. Overall, tools that used only long-read data performed well across datasets, suggesting a paired short-read dataset is not necessary anymore. We observed that method performance is dependent on sequencing depth and sequencing quality. For instance, except snuupy which failed to process PromethION data, most tools perform better with PromethION data than MinION data, and with the R10.4.1 rather than with the R9.4.1 chemistry. The performance also depends on the task. For example, wf-single-cell performs better in the gene expression quantification, but is surpassed by Bambu and Isosceles at isoform discovery. In contrast Bambu misidentifies UMI sequences leading to an overestimation of RNA counts.

Unexpectedly, long-read methods, particularly Bambu and wf-single-cell, outperform CellRanger in the characterization and annotation of cell types. This is likely due to the 3’ bias in short-read data, which can compromise accurate gene expression detection. In line with this, Bambu and wf-single-cell consistently detect a greater number of genes per cell than CellRanger. Similarly, when assessing concordance with short-read data, a discrepancy emerges: tools that show the highest correlation with short-read datasets do not necessarily perform best on simulated data. This further supports recent findings (53–55) showing that short-read data are not free from bias, reinforcing the value of long-read approaches for transcriptomic profiling.

As expected, we observed a consistent relation between UMI error correction and the accuracy of gene and transcript expression quantification. For instance, wf-single-cell employs an advanced UMI error correction strategy based on a directed graph algorithm, resulting in the highest fidelity in gene/transcript expression estimation. In contrast, Bambu does not perform UMI error correction, which leads to erroneous UMI deduplication and consequently an overestimation of expression counts—particularly under conditions of high sequencing saturation and high sequencing error rates. Notably, this overestimation strongly correlates with the expected expression levels.

The higher sequencing error rate observed in long-read data, compared to short-read sequencing, adversely affects the accuracy of read alignment, particularly at splice junctions (56). To overcome this limitation, error correction approaches such as consensus sequence generation, as implemented in Sicelore, or splice alignment correction, as adopted in Bambu, can be applied (57). Consistent with this, our results showed that Sicelore 2.1 achieves more accurate read assignment to annotated transcript models, while Bambu provides the most reliable transcript assembly under high-error condition. Moreover, coupling wf-single-cell with Isosceles or IsoQuant tends to predict more false-positive novel transcript models, while coupling Sicelore 2.1 with Isosceles or IsoQuant results in splice alignment errors.

Despite the rapid development of single-cell and spatial long-read RNA-sequencing methods, setting up unbiased and objective benchmarking remains challenging. A major barrier to systematic benchmarking is the lack of standard output formats. Specifically, we faced issues to automate the parsing of the count matrix. Indeed, depending on the tools, matrices are either stored in sparse or dense format, with or without the presence of metadata, such as exon sizes. Overcoming the technical challenges of output formats would enable the introduction of continuous and automated comparisons across tools.

In contrast to bulk and single-cell short-read RNA-seq, for which extensive public data for benchmarking are available (53, 58–60), single-cell and spatial long-read transcriptomics still lacks comprehensive benchmark resources. This represents a substantial obstacle to optimize long-read methods for transcriptome profiling. Although long-read RNA-seq datasets are increasingly available for bulk experiments (38, 55, 57), resources for single-cell and spatial contexts remain limited. In this study, we aimed to address this gap by providing access to a collection of datasets specifically generated across different sequencing protocols, depths, and chemistries. However, some aspects of data variability were not captured, for example, other long-read platforms such as PacBio or additional library types like 10x Genomics 5’ kits, which are supported by several of the tools evaluated in this study. Of note, a recent study suggests that PacBio outperforms Nanopore in terms of cell type annotation, novel isoforms identification and allele-specific expression (13).

One of the key advantages of long-read RNA-seq is its ability to generate full-length reads, enabling the identification of complete fusion transcripts or identify variants in single-cell and spatial contexts. Despite the importance of these features, our benchmark focused on core transcriptomic tasks, such as UMI/barcode detection, gene and isoform quantification, clustering, and annotation due to the current lack of standardized tool outputs and methods for evaluating fusion detection and variant calling in these settings.

Altogether, this study offers three key contributions: (i) a collection of single-cell long-read datasets specifically designed to support the benchmarking of bioinformatics tools, (ii) an independent evaluation of ten computational methods providing a practical resource to select the most appropriate tools according to the desired analysis, and (iii) a reusable benchmarking workflow enabling method developers to compare their own approaches against the set of reference methods evaluated in this work.

## Materials and Methods

### Data

#### Single-cell cDNA library preparation and sequencing. Generation of single-cell cDNA

The MPNST1 and MPNST2 single-cell suspension were converted into a barcoded scRNA-seq library with the 10x Genomics Chromium Single Cell 3’ Library, Gel Bead & Multiplex Kit and Chip Kit (v3), aiming for 20,000 cells following the manufacturer’s instructions. The following modifications were applied to MPNST1: we extended the PCR elongation time during the initial PCR amplification of the cDNA from the manufacturer recommended 1 min to 3 min to minimize preferential amplification of small cDNAs (16).

#### Short-read Illumina libraries

Half of the amplified cDNA was used for short-read sequencing library preparation following the 10x Genomics protocol. MPNST1 was sequenced on an Illumina Nextseq 500 sequencer at GenomiqueENS facility, resulting in 491 M reads. MPNST2 was sequenced on an Illumina Nextseq 2000 sequencer at IMRB facility, resulting in 401 M reads.

#### Long-read ScNaUmi-seq protocol libraries

MPNST1 and MPNST2 libraries were prepared as described in Lebrigand et al. (16), including the Optional steps for the depletion of cDNA lacking a terminal poly(A)/poly(T) tail. Nanopore sequencing libraries were prepared with the Oxford Nanopore SQK-LSK-110[MOU2] kit following the manufacturer’s instructions. Sequencing was performed on R9.4.1 flow cells at GenomiqueENS facility (MPNST1 on MinION, MPNST2 on a P2solo) and at the Centre National de Recherche en Génomique Humaine (CNRGH) facility (MPNST1 PromethION).

#### Single-cell long-read ONT protocol libraries

The MPNST2 library was prepared with the “Single Cell sequencing on Prome-thion protocol” (https://nanoporetech.com/document/single-cell-transcriptomics-with-cdna-prepared-using-10x) with the SQK-PCS11 kit. We re-amplified 10 ng of the 10x Genomics PCR product for 4 cycles with 5’-CAGCTTTCTGTTGGTGCTGATATTGCAAGCAGTGGTATCAACGCAGAG-3’ and 5’ Biotine-CAGACACTTGCCTG TCGCTCTATCTTCCTACACGACGCTCTTCCGATCT-3’. After 0,8x AmpureXP purification to remove excess biotinylated primers, biotinylated is bound to Dynabeads™ M-280 Streptavidin beads (Invitrogen) and amplified with the primers cPRM for 4 cycles. Amplified cDNA was purified with 0,8x Ampure XP and sequenced on a R9.4.1 flowcell at GenomiqueENS facility on a P2solo according to the manufacturer’s protocol.

#### Spatial scRNAseq long-read data

Spatial long-read data were downloaded from Gene Expression Omnibus (accession number: GSE153859), both as FASTQ and count matrix, according to the benchmarking steps. The subsequent processing was performed with the same pipeline as the non-spatial data.

#### Simulation of scRNAseq long-read data

To simulate scRNAseq Nanopore datasets (Table S1), the FASTQ file from the MPNST1 ScNaUmi-seq PromethION dataset was processed with wf-single-cell v3.0.3 using default parameters. The raw transcript count matrix was then analyzed using the Seurat package (v4.3.0). Normalization was performed using the NormalizeData function with the LogNormalize method, followed by log-transformation and scaling using ScaleData. A total of 2,000 variable features were selected, and 15 principal components were retained for dimensionality reduction. Clustering was performed using the Louvain algorithm with a resolution of 0.5, and cell type annotation was based on a predefined set of marker genes (defailed below). The transcript count matrix was then filtered to retain only macrophages, fibroblasts, and endothelial cells based on the annotation results. The filtered matrix was then used as input for AsaruSim v1.0.3 (42).

To generate realistic reads and replicate the error profile from the MPNST1 ScNaUmi-seq PromethION dataset (R9.4.1 Nanopore chemistry), a subset of the real FASTQ file was used as a reference model to simulate sequencing errors, and quality scores. The same subset of the real FASTQ file was aligned to the reference transcriptome using minimap2 (61) to derive an empirical truncation probability distributions on both 5’ and 3’ ends. A total of 39,816 transcripts were simulated across 635 cells. Subsequently, 20 million reads were randomly selected from the simulated pool following an 8-cycle PCR amplification. Additional FASTQ files were generated using the same procedure, but employing a real FASTQ file derived from R10.4.1 Nanopore chemistry as the reference model.

To simulate multiple datasets with various UMI duplication rates, we first analyzed the MPNST1 ScNaUmi-seq Illumina dataset and identified 4,500 cell-associated barcodes using Cell Ranger v6.0.0. We then used the corresponding UMI counts as input for AsaruSim v1.0.3 (–CB_counts option) to simulate long-read data, ensuring a realistic read count distribution. Then, a total of 10,000 cDNA sequences were subsampled from the Ensembl release-113 mouse transcript reference with seqtk (62) using a random seed value of 123 and used as reference transcriptome for AsaruSim. Five datasets were generated with increasing UMI duplication levels (–umi_duplication option) ranging from 1 to 5. Finally, 1 million reads were randomly selected from each simulated dataset, achieving sequencing saturation levels of 0.20, 0.35, 0.45, 0.54, and 0.60, respectively.

### Time and memory

To measure execution time and peak memory usage, reported in Figure S1, all jobs were executed on Linux servers equipped with Intel Xeon processors, 32 CPU cores, and 190 GB of memory. A maximum of 20 threads was allocated for all jobs. Execution time and maximum memory usage were assessed using the Linux time (/usr/bin/time) command with the -v flag. Execution time was extracted from the “Elapsed time” field, while peak memory consumption was recorded from the “Maximum resident set size” field.

### Data processing

#### Definitions

We call barcode the 16-nucleotide sequence used to uniquely identify droplets or spots. All the possible sequences are referenced in an inclusion list (formerly called whitelist), provided by 10X Genomics. After sequencing, barcodes are generally compared to the inclusion list to discard unreferenced sequences. The sequenced barcodes found in the inclusion list correspond to the barcode shortlist. This shortlist is then further filtered to remove empty droplets/spots. A knee plot, depicting the number of UMI associated with each barcode from the shortlist, enables distinguishing empty from non-empty droplets. The barcodes associated with non-empty droplets are finally called cell-associated barcodes.

#### Tools and parameters associated with short-read data

For short-read (Illumina) MPNST datasets, BCL files were processed using 10x Genomics Cell Ranger software (v3.0.0). Reads were mapped onto a custom mouse transcriptome based on GRCm38 transcriptome, in which we added tdTomato sequence contained in the tumor cells. For Nanopore data Fast5 files, base calling was performed with Guppy v4.2.2 (https://nanoporetech.com/software/other/guppy), followed by read quality control using ToulligQC v2.3 (https://github.com/GenomiqueENS/toulligQC). The sequencing saturation curves were computed using CellRanger v6.0.0 for short-read data and wf-single-cell v3.0.1 for Nanopore data. The numerical values are shown in Table S1.

#### Tools and parameters associated with long-read data

The wf-single-cell pipeline was executed with recommended parameters and the –expected_cells option set to 900 for spatial datasets, 4,500 for the MPNST1 dataset, and 11,000 for the MPNST2 dataset. In addition, the –barcode_max_ed parameter was set to 1 to improve barcode assignment accuracy.

For Isosceles, the tool was run using the recommended settings in two separate workflows: (1) with the deduplicated BAM file generated from step 4b (consensus sequences per UMI) of the Sicelore v2.1 Nextflow workflow, and (2) with the tagged BAM output of wf-single-cell. For the second workflow, the bam_to_tcc function was applied using the parameter barcode_tag=“CB”. The tagged BAM files were deduplicated using the “dedup” command from UMI-tools (43). In both cases, the GTF annotation file provided by Isosceles contained only exonic features and was fixed to add transcripts using AGAT’s script agat_convert_sp_gxf2gxf.pl (63).

scNanoGPS was run with default parameters, except for – CB_mrg_thr 1 and –CB_no_ext 0.3, which were adjusted to enhance barcode shortlist validation for PromethION and simulated datasets. For the simulated amplified datasets, the –min_gene_no parameter was set to 10 to prevent overly stringent filtering of gene count matrices.

FLAMES was executed by first running the precompiled C++ binary src/bin/match_cell_barcode with an edit distance of 2, a 12 nucleotide UMI length, and the CellRanger inclusion list (formerly whitelist) as input. Subsequently, the sc_long_pipeline.py script was executed with default parameters.

BLAZE was executed using –expected_cells set to 900 for spatial data, 4,500 for MPNST1, and 11,000 for MPNST2. The run included the following additional options: –high-sensitivity-mode, –emptydrop, and –out-CB-whitelist=whitelist_hs.

Snuupy and Sicelore were executed with recommended parameters. For both tools, barcode shortlisting was based on the CellRanger-provided inclusion list, using an edit distance of 2 for both barcode and UMI matching. Sicelore v2.1 was also run independently with default parameters, without supplying an inclusion list.

To run the long-read-only tools with spatial transcriptomics datasets, the barcode inclusion list visium-v1.txt was used in place of 3M-february-2018.txt, following guidance from 10x Genomics (https://kb.10xgenomics.com/hc/en-us/articles/115004506263-What-is-a-barcode-inclusion-list-formerly-barcode-whitelist).

Under these conditions, all tools received as input the FASTQ files, reference transcriptome sequence and annotation, and output a cells-by-isoforms count matrix. For some tools (detailed below), isoform discovery is performed and the output additionally includes the new annotation file. All software environments and version details are provided in Table S3.

### Accuracy in barcodes and UMI assignment

#### Evaluation of barcode false-positive detection

To assess the false-positive detection rate in barcode calling for each method in Figure 4b, we focused on tools that identify cell-associated barcodes based on read count thresholds: BLAZE, wf-single-cell, scNanoGPS, and Bambu. These methods detect barcodes by counting their occurrences and selecting the most abundant ones as cell-associated (shortlisted) barcodes, typically using a threshold determined by the inflection point in the knee plot of barcode rank versus UMI count. To evaluate performance, we computed precision-recall curves across varying count thresholds, using the shortlist of barcodes identified by CellRanger on short-read data (raw count matrix) as the ground truth. Prior to this, empty droplets were filtered out using the DropletUtils R package (45), with default parameters and a false discovery rate (FDR) threshold of 1%. Barcodes identified in long-read data but not present in the CellRanger-filtered shortlist were considered false positives, while true positives were defined as barcodes present in both sets.

#### Accuracy of UMI error correction

To generate Figure 4c-d, for each tool, UMI sequences were extracted from tagged BAM files prior to the deduplication step using bioalcidaejdk (64). Raw and corrected UMI sequences were obtained as follows: for Sicelore 2.1, the ‘U7’ tag was used for raw UMIs and the ‘U8’ tag for corrected UMIs; for wf-single-cell, the ‘UR’ and ‘UB’ tags denoted raw and corrected UMIs, respectively. In the case of FLAMES, raw UMIs were parsed from read names in the realign2transcript.bam file, while corrected UMIs were extracted using a custom script following the UMI merging step. For scNanoGPS, raw UMI sequences were retrieved from the barcode_list.tsv file, and corrected UMIs were extracted from read names after the ‘curation’ step. For Bambu, raw UMIs were obtained from the demultiplexed.bam file.

To evaluate UMI correction accuracy, we compared the corrected and raw UMIs against the ground-truth UMI sequence from simulated data. Precision, recall, and F1-score were calculated using the following definitions. A true positive (TP) was defined as a corrected UMI that matched the ground-truth UMI. Precision was computed as the proportion of correctly corrected UMIs out of all corrected UMIs (TP / total corrected), while recall was the proportion of correctly corrected UMIs out of all ground-truth UMIs (TP / total true). The F1-score was computed as the harmonic mean of precision and recall:

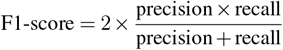

In addition, clustering accuracy of UMI grouping was assessed using the Adjusted Rand Index (ARI), computed between the true UMI group labels and the predicted UMI clusters provided by each tool. All analyses were implemented in R using traceable R Markdown notebooks.

### Count matrix processing

#### Projection and cell type annotation

The 10x Cell Ranger filtered matrix output files were imported into Seurat (v4.3.0) (65). For all samples and for quality control purposes, genes expressed in less than three cells were discarded, and cells with less than 300 unique gene counts were filtered out. The gene expression matrix was normalized with Seurat’s LogNormalize function and 2,000 variable features were selected using Seurat’s vst method then the matrix was scaled using Seurat’s linear model. A PCA was generated using Seurat’s RunPCA function. Then, cells were clustered using the Louvain algorithm, with 15 principal components as input of Seurat’s FindNeighbors function and a resolution of 0.5 for Seurat’s FindClusters function. Single cells were annotated for cell type using a modified version of Seurat’s AddModuleScore function and cell type-specific marker gene sets (see Code and Data availability).

#### Pseudo-bulk expression correlation analysis

Pseudo-bulk analyses were used for Figure 4g, Figure 7c and Figure S2e-f. To compare pseudo-bulk expression profiles between methods, Seurat objects were annotated for cell types. Gene counts were aggregated across cells within each cell type to generate pseudo-bulk matrices. These matrices were normalized using counts-per-million followed by log-transformation. Pearson correlation coefficients were then computed between matched cell types across methods.

#### Quantitative assessment of cell type annotation and projection

To evaluate the biological resolution achieved by each method, we assessed the quality of cell type annotation and projection. The results are presented in Figure 5 and Figure 6.

First, individual cells were annotated based on specific gene markers, following the annotation protocol described below. We computed the Local Inverse Simpson’s Index (LISI) (46), a diversity score designed to assess both batch mixing (iLISI) and cell-type separation (cLISI). The cLISI score represents the effective number of cell type labels present in the neighborhood of each cell on the UMAP projection. cLISI scores were computed for each method using the compute_lisi function from the lisi R package, applied to the UMAP reduction of cell type–annotated Seurat objects.

Next, the long-read gene count matrix was integrated with the short-read count matrix using Seurat’s CCAIntegration method, after which cLISI scores were computed again using the same compute_lisi function. To assess the similarity between cell type labels derived from short-read and long-read data, we used standard clustering evaluation metrics, including the Adjusted Rand Index (ARI).

For spatial transcriptomics datasets, LISI was adapted to quantify the effectiveness of spatial domain detection. In this context, the LISI score reflects the effective number of spatial domain labels present within the local neighborhood of each spot, using a fixed 5-spot window. In addition to LISI and ARI, we employed the CHAOS score to evaluate the spatial continuity of the detected domains (66, 67). The CHAOS score quantifies intra-domain spatial compactness by calculating the average distance to the nearest neighbor within each domain, normalized by the total number of spots. Lower CHAOS values indicate more spatially coherent clusters.

#### Differential isoform analysis

Differential isoform analysis was performed using the Isoswitch R package (https://github.com/ucagenomix/isoswitch). Results are presented in Figure 7d and Figure S4. For each tool, gene- and transcript-level count matrices were used to construct a multi-assay Seurat object. Each assay was normalized independently using the LogNormalize method. Clustering was performed based on the gene-level assay, and ground-truth cell type annotations were subsequently imported.

Differential isoform usage was then assessed using the ISO_SWITCH_ALL function from Isoswitch, which lever-ages Seurat’s FindMarkers function to identify differentially spliced genes (DSGs) for each cell type (macrophages, fibroblasts, and endothelial cells). The resulting lists of DSGs were used to generate an UpSet plot to visualize overlaps across tools (Figure S4). In addition, log fold changes (logFC) were computed and correlated with those from the ground truth to assess consistency in differential isoform detection.

#### Gene ontology analysis

DSGs were filtered for p-value < 0.01 and absolute value of the average log2FC > 2. For gene ontology analysis, the list of 2,942 significant DSGs was provided as input (gene names) to the clusterProfiler’s enrichGO function (version 4.6.2) using default parameters (68). Top 8 ontologies were visualized using enrichplot (version 1.18.4).

### Genes and isoforms identification and quantification

We used the simulated data to assess the accuracy of transcript assignment.

#### Accuracy of reads assignment to isoforms

Transcript labels were extracted from tagged BAM files using bioalcidaejdk (64). The ‘TR’ tag was used for Sicelore 2.1, and the ‘IT’ tag for wf-single-cell. For FLAMES, transcript labels were parsed from the realign2transcript.bam file. For scNanoGPS, a customized version of LIQA (69) was used to extract read names and their assigned transcript labels. To evaluate transcript assignment accuracy, assigned transcript labels were compared to the simulated ground-truth transcript IDs. For each transcript class, precision and recall were computed using parallel processing across all simulation cycles. The F1-score was then calculated as the harmonic mean of precision and recall. Precision, recall, and F1-score values were computed per tool for 1 and 5 UMI duplication simulation, and summarized as mean ± standard deviation for each metric. Results are shown on Figure 7a.

#### Comparison of gene and isoform expression estimation to ground truth

To assess the accuracy of gene and isoform expression quantification in Figure 7b, we compared the output of each method to the simulated ground truth using the root of mean squared error (RMS error, RMSE):

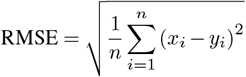

where *x*_*i*_ and *y*_*i*_ represent the estimated and true expression values for gene *i*, respectively.

To control for variation in sequencing depth and mitigate library size biases, each expression matrix (estimated and ground truth) was standardized using the mean and standard deviation derived from the ground truth. After normalization, the RMSE was calculated between the normalized matrices to provide a more robust comparison of expression accuracy at the single-cell level. All evaluations were performed separately for each simulation and method. Results are shown on Figure 4e-f and Figure S2d-e.

### Novel isoform prediction

Isoform prediction, or transcriptome re-annotation, is a feature integrated in some tools (Bambu, IsoQuant, FLAMES, Isosceles and wf-single-cell). We evaluated the output annotation files as follows.

#### Discovery of novel isoforms in simulated data

To evaluate the accuracy of novel transcript prediction, we generated a reduced reference annotation by excluding 15% (4,412 transcripts) of the 29,413 expressed transcripts from the GENCODE M35 annotation used in the simulated dataset. The excluded transcripts were treated as the true novel isoforms, while the remaining annotated transcripts were considered as true known isoforms.

The incomplete reference annotation, along with the full set of simulated reads, was then provided to isoform prediction tools: wf-single-cell, FLAMES, and Bambu. For Isosceles, raw reads were first processed using Sicelore v2.1 or wf-single-cell then tagged and deduplicated BAM files were used as input. The resulting transcriptome assemblies, output in GTF format, were then parsed to classify transcripts as either known or novel based on the annotation status in the GTF files.

To assess accuracy, we used gffcompare to compare the predicted novel transcripts against the set of true novel isoforms. Precision and sensitivity were computed, and the F1-score was calculated as the harmonic mean of precision and sensitivity. Results are shown on Figure 7e.

#### Consistency among predicted annotations on real data

To evaluate the overlapping between annotations, we used gffcompare to pairwise compare the full annotation which was predicted by each tool (predicted_annotations.txt lists all the annotation files) and the original annotation (ref_annotation.gff). We used the command gffcompare -g ref_annotation.gff -i predicted_annotations.txt. This command outputs isoform class codes that were converted to SQANTI classification (70). The results are visualized using an upset plot on Figure 7f.

## Code and Data Availability

Sequencing data related to the MPNST1 and MPNST2 samples are available on ArrayExpress, under accession number E-MTAB-15190. It includes the FASTQ files and the count matrices obtained with the wf-single-cell pipeline. The workflow behind the benchmark, called scKeñver, is available on Github (https://github.com/alihamraoui/scKenver). The specific version used here is available on Zenodo (record ID: 16098020). This repository further contains the data simulated with AsaruSim and the gene markers used for cell type annotation.

## Author Contributions

Conceptualization: AH, MTC; Software: AH, AO; Data curation: AH, AO; Formal analysis: AH, SL; Investigation: AH, AO, CSB, FC, SL; Methodology: AH, AO, MTC; Visualization: AH; Resources: CSB, FC; Supervision: SL, LJ, MTC; Funding acquisition: MTC; Project administration: MTC; Writing – original draft: AH, AO, MTC; Writing – review & editing: AH, AO, SL, LJ, MTC.

## Acknowledgments

We thank Alice Lebreton, Stéphane Le Crom and Piotr Topilko for insightful discussions regarding this work.

## Funding Sources

The GenomiqueENS core facility was supported by the France Génomique national infrastructure, funded as part of the “Investissements d’Avenir” program managed by the Agence Nationale de la Recherche (contract ANR-10-INBS-09). This work was conducted with financial support from ITMO Cancer of Aviesan on funds administered by Inserm (contract C21049DS). A CC-BY public copyright license has been applied by the authors to the present document, in accordance with the grant’s open access conditions.

## Supplementary Figures

**Supplementary Figure 1.**
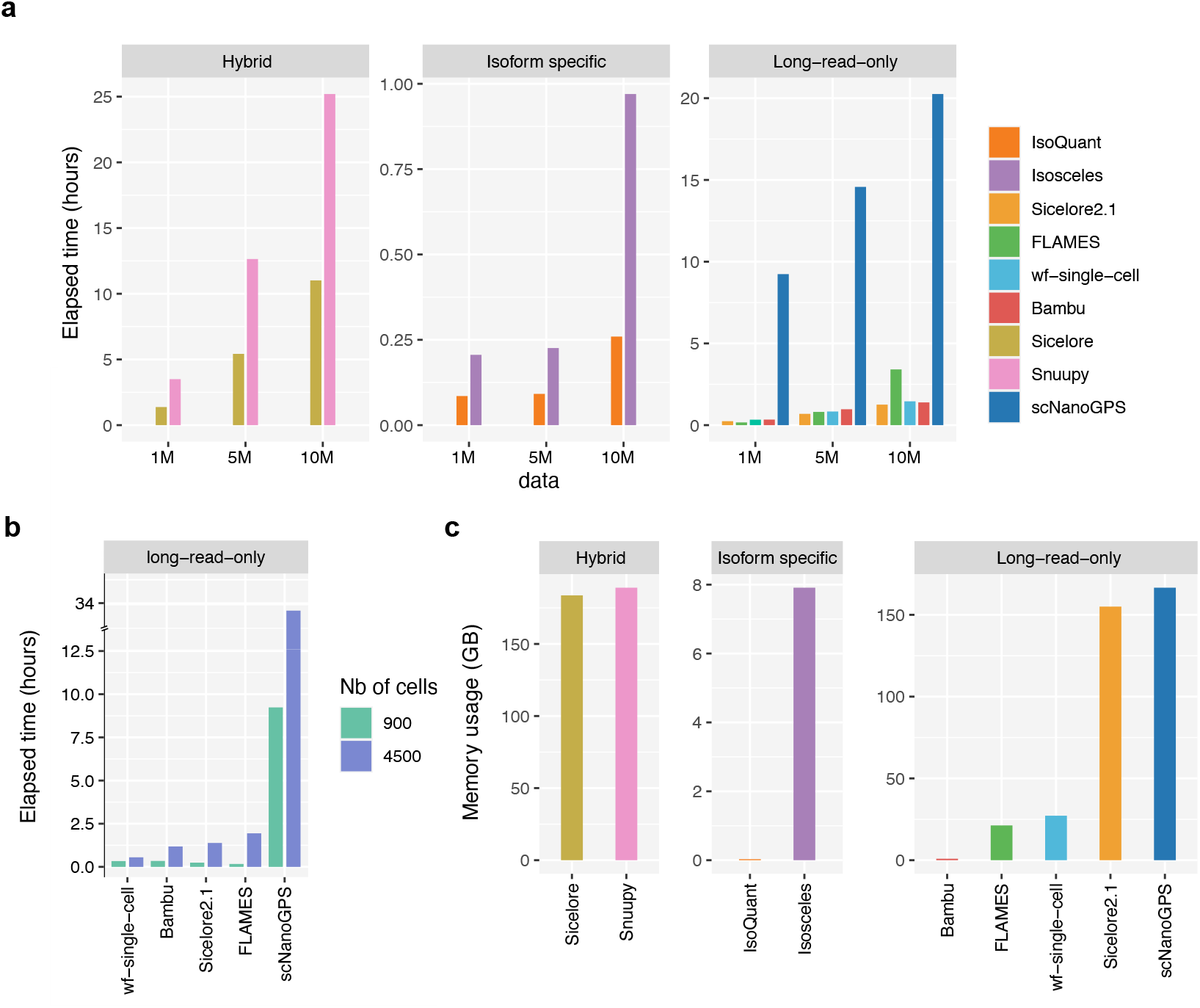
Running time and memory usage —. **(a)** Elapsed time in hours for hybrid, long-read-only and isoform specific methods computed for 1, 5 and 10 million reads obtained by subsampling the MOB datasets. scNapBar is not included due to exceeding running time. **(b)** Elapsed time in hours for long-read-only methods computed for 1 million reads from 900 cells of the MOB datasets versus 1 million reads from 4,500 cells in simulated datasets. **(c)** Peak of memory used by hybrid, long-read-only and isoform specific methods used for 10 millions reads from the MOB datasets.

**Supplementary Figure 2.**
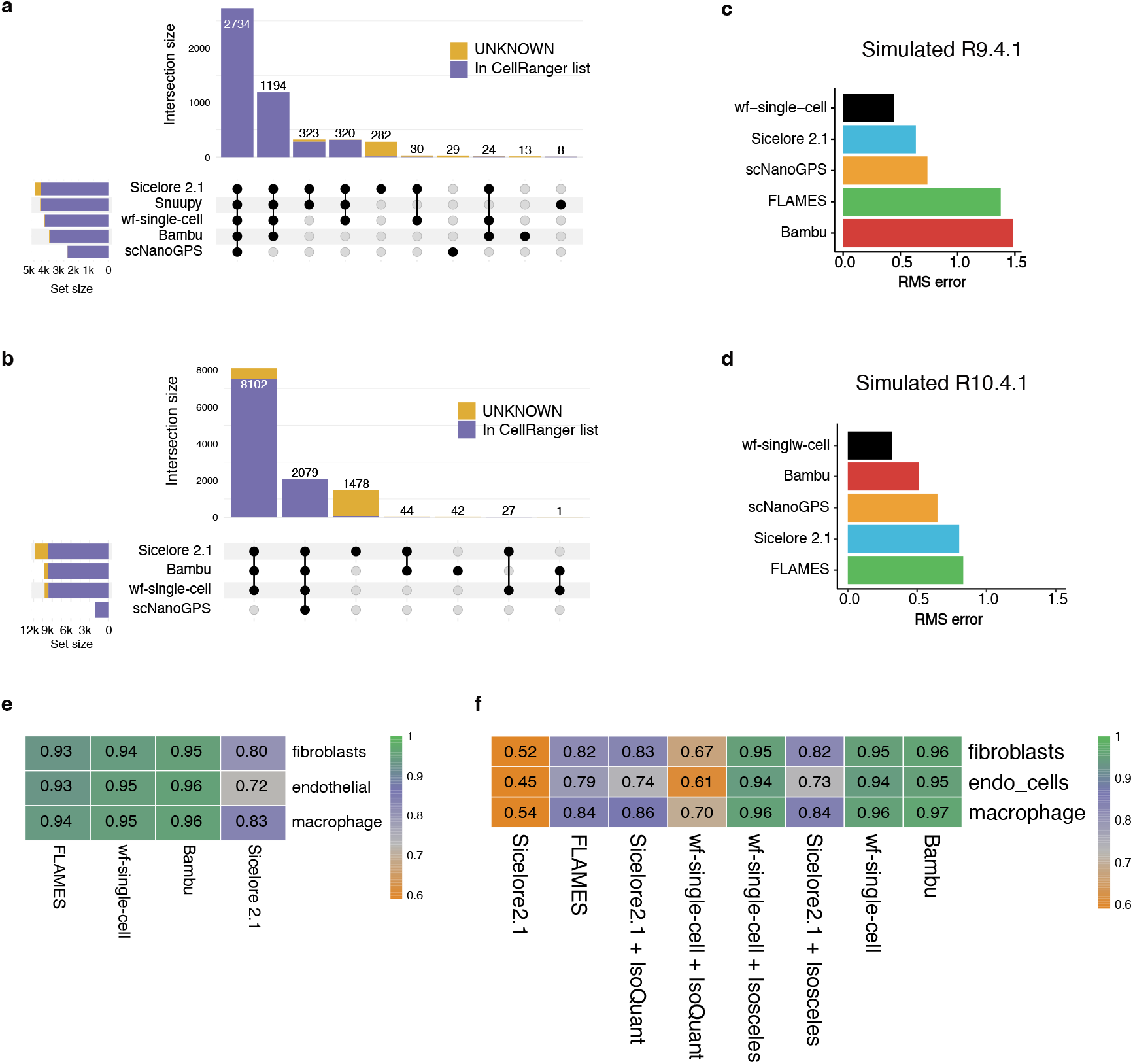
Accuracy in barcode assignment and gene quantification fidelity —. **(a)** Barcode upset plot comparing different shortlists. The bar chart on the left shows the total number of barcodes found by each tool. The bar chart on top shows the number of barcodes in the intersection of shortlists from specific combinations of methods. The dots and lines underneath show the combinations. **(b)** The same as plot a for MPNST ONT data. **(c-d)** RMSE between the observed and expected gene expression values. RMSE were computed following normalization, where each expression matrix was standardized using the mean and standard deviation of the ground truth in the Simulated R9.4.1 **(c)** or R10.4.1 **(d)** data. **(e)** Pearson’s correlation (color scale) of each method’s gene quantifications with expected quantification at pseudo-bulk level in R10.4.1 data. **(f)** Pearson’s correlation (color scale) of each method’s isoform quantifications with expected quantification at pseudo-bulk level in R10.4.1 data.

**Supplementary Figure 3.**
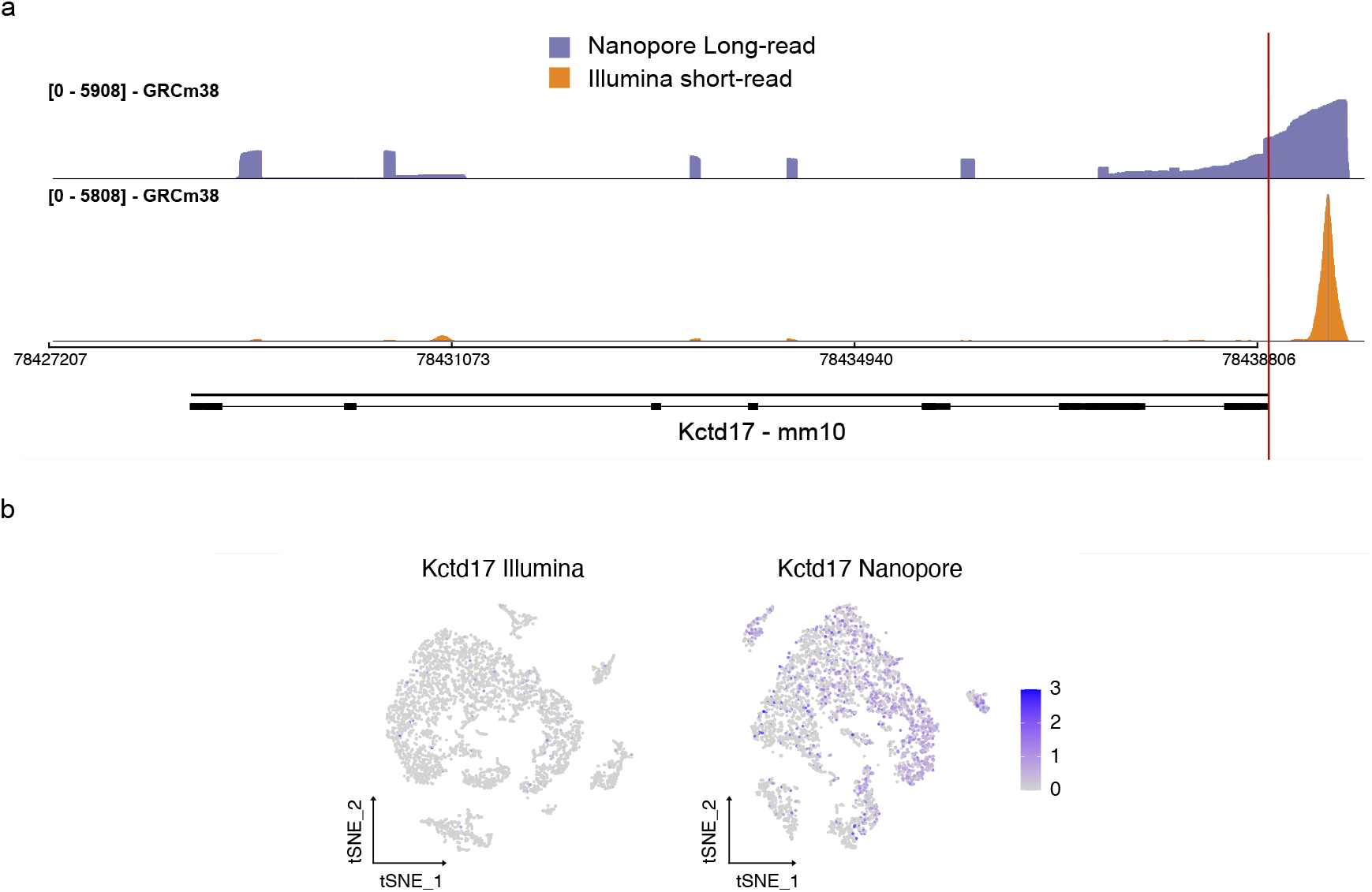
Importance of gene annotation for isoform detection —. **(a)** Sashimi plot showing Illumina short reads and Nanopore long reads aligned to GRCm38 (mm10) mouse genome. Due to incomplete 3’ UTR annotation of *Kctd17* in ensemble annotation v86, a large fraction of short reads map to non-genic regions, leading to quantification errors in short-read data. **(b)** FeaturePlot of *Kctd17* gene in MPNST1 Illumina and PromethION data.

**Supplementary Figure 4.**
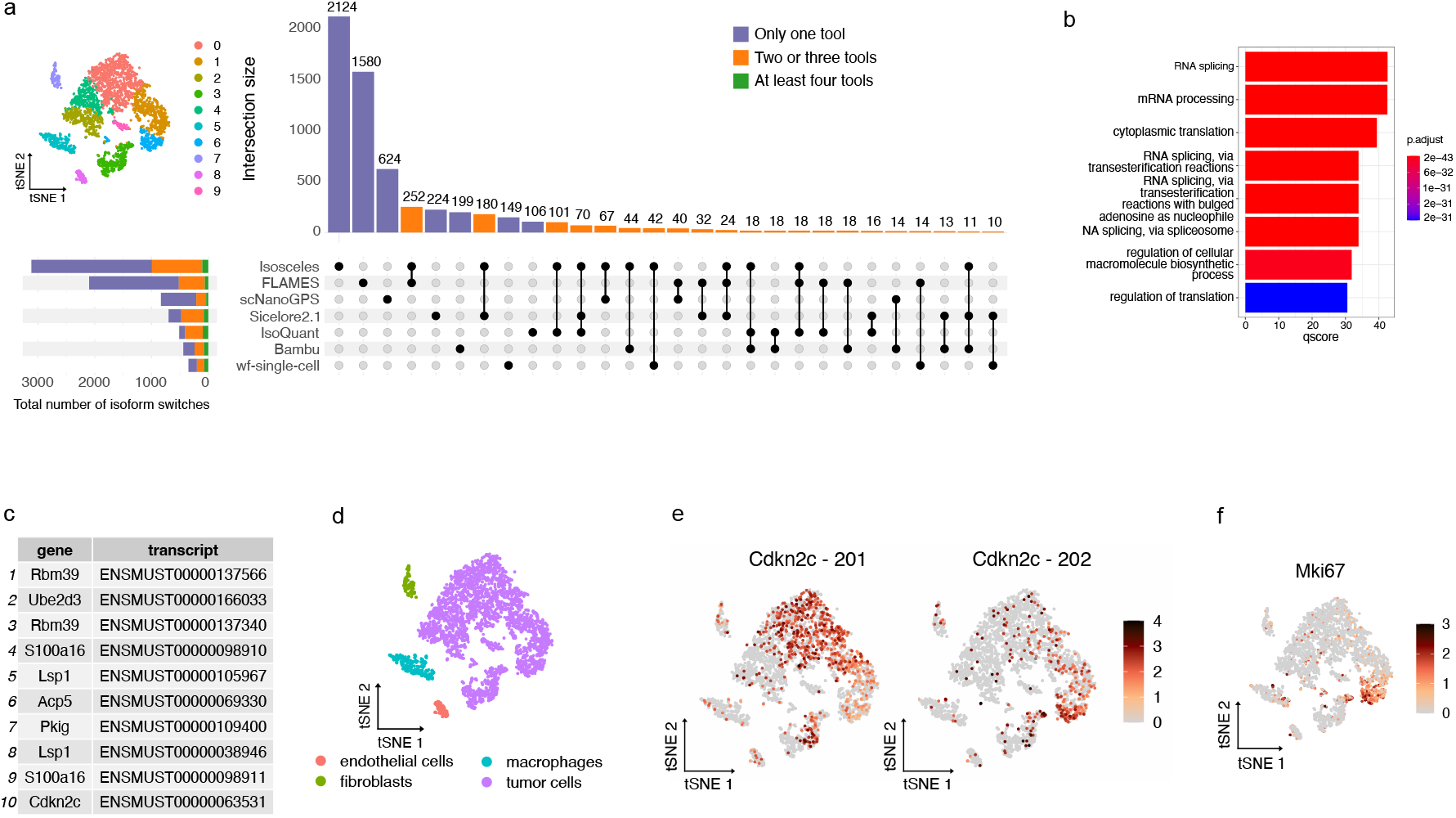
Comparison of the genes showing an isoform switch, depending on the preprocessing tools —. **(a)** (left) tSNE projection of the MPNST1 PromethION dataset at the gene level. Cells are colored according to 10 cell clusters. (right) Upset plot comparing the number of isoforms characterized by a switch between any two cell clusters, for each preprocessing tool. The bars are colored according to the number of tools having identified the isoforms. In total, 6,266 isoforms were identified characterized by a switch, corresponding to 2,942 genes. **(b)** Barplot showing the top 8 enriched gene ontologies associated with the 2,942 genes characterized by an isoform switch. **(c)** Top 10 genes showing an isoform switch between any two cell clusters, displayed with their Ensembl transcript identifier. **(d)** tSNE plot with cells colored according to four cell types. **(e)** tSNE plot depicting the isoform switch of *Cdkn2c* gene, with differential usage of the isoforms 201 and 202 of this gene. **(f)** tSNE plot depicting the expression of the *Mki67* gene, used as a proxy for proliferative activity.

## Supplementary Tables

**Supplementary Table 1.** Datasets used in this study — -: not available

| Sample | Sequencing platform | Sequencing protocol | Sequencing kit | Sequencing saturation % | No. cells/spots | Total reads | Median phred score |
| --- | --- | --- | --- | --- | --- | --- | --- |
| MPNST1 | ONT PromethION | scNaUmi-seq | SQK-LSK110 | 35% | 4,559 | 151,334,327 | 11.32 |
| MPNST1 | ONT MinION | scNaUmi-seq | SQK-LSK109 | 4% | 4,559 | 9,281,420 | 11.75 |
| MPNST1 | Illumina NextSeq 500 | 10x Genomics | - | 81% | 4,559 | 491,161,783 | - |
| MPNST2 | ONT PromethION | scNaUmi-seq | SQK-LSK110 | 8% | 10,297 | 130,507,091 | 14.0 |
| MPNST2 | ONT PromethION | ONT | SQK-PCS111 | 6.7% | 10,297 | 108,009,263 | 13.56 |
| MPNST2 | Illumina NextSeq 500 | 10x Genomics | - | 34% | 10,297 | 401,224,345 | - |
| MOB (Lebrigand et al 2023) | Illumina NextSeq 500 | 10x Genomics | - | 93.1% | 900 | 253,954,360 | - |
| MOB (Lebrigand et al 2023) | ONT PromethION | scNaUmi-seq | SQK-LSK-109 | 57% | 900 | 10,226,285 | 11.81 |
| Simulated R9.4.1 | simulation | - | - | 79% | 635 | 20,000,707 | 13 |
| Simulated R10.4.1 | simulation | - | - | 86% | 635 | 20,000,261 | 20 |
| Simulated dup1 R9.4.1 | simulation | - | - | 20% | 4,559 | 990,690 | 13 |
| Simulated dup2 R9.4.1 | simulation | - | - | 35% | 4,559 | 1,006,813 | 13 |
| Simulated dup3 R9.4.1 | simulation | - | - | 45% | 4,559 | 991,685 | 13 |
| Simulated dup4 R9.4.1 | simulation | - | - | 54% | 4,559 | 995,366 | 13 |
| Simulated dup5 R9.4.1 | simulation | - | - | 60% | 4,559 | 991,390 | 13 |
| Simulated dup1 R10.4.1 | simulation | - | - | 20% | 4,559 | 990,690 | 20 |
| Simulated dup2 R10.4.1 | simulation | - | - | 35% | 4,559 | 1,006,813 | 20 |
| Simulated dup3 R10.4.1 | simulation | - | - | 45% | 4,559 | 991,685 | 20 |
| Simulated dup4 R10.4.1 | simulation | - | - | 54% | 4,559 | 995,366 | 20 |
| Simulated dup5 R10.4.1 | simulation | - | - | 60% | 4,559 | 991,390 | 20 |

**Supplementary Table 2.** scRNA-seq long-read methods evaluated in this study — *: paired dataset sequenced in short-read is used by hybrid approaches. References: 1 (Lebrigand et al.), 2 (Long et al.), 3 (Wang et al.), 4 (Tian et al.), 5 (https://github.com/epi2me-labs/wf-single-cell), 6 (Shiau et al.), 7 (You et al.), 8 (Sim et al.), 9 (Kabza et al.), 10 (Prjibelski et al.).

| Tool name | Programming language | Barcode demultiplexing | Chimeric reads filtering | UMI error correction | UMI deduplication | Isoform analysis | Transcript discovery |
| --- | --- | --- | --- | --- | --- | --- | --- |
| Sicelore<sup>1</sup> | Java | short-read data | yes | SR data | consensus sequence | yes | — |
| Snuupy<sup>2</sup> | Python | short-read data | yes | SR data | consensus sequence | PolyA-based | — |
| scNapBar<sup>3</sup> | C++/Python/Perl | short-read data | — | SR data | — | yes | — |
| FLAMES<sup>4</sup> | C++/Python | — | — | Merge UMIs within a fixed edit distance | keep longest read in each UMI group | yes | yes |
| wf-single-cell<sup>7</sup> | Python | 10X inclusion list (count over threshold) | yes | UMI-tools<sup>5</sup> clustering algorithm | UMI-tools<sup>5</sup> representative | StringTie2<sup>6</sup> / FLAMES | yes |
| Sicelore 2.1<sup>1</sup> | Java | 10X inclusion list | yes | Cluster UMIs based on edit distance | consensus sequence | yes | — |
| scNanoGPS<sup>9</sup> | Python | no guidance (count over threshold) | yes | Merge UMIs within a fixed edit distance | consensus sequence | LIQA<sup>8</sup> | — |
| BLAZE<sup>10</sup> | Python | 10X inclusion list (count over threshold) | — | — | — | — | — |
| Bambu<sup>11</sup> | R | 10X inclusion list (count over threshold) | yes | — | keep longest read in each UMI group | EM algorithm | yes |
| Isosceles<sup>12</sup> | R | — | — | — | — | EM algorithm | yes |
| IsoQuant<sup>13</sup> | Python | — | — | — | — | EM algorithm | yes |

**Supplementary Table 3.**
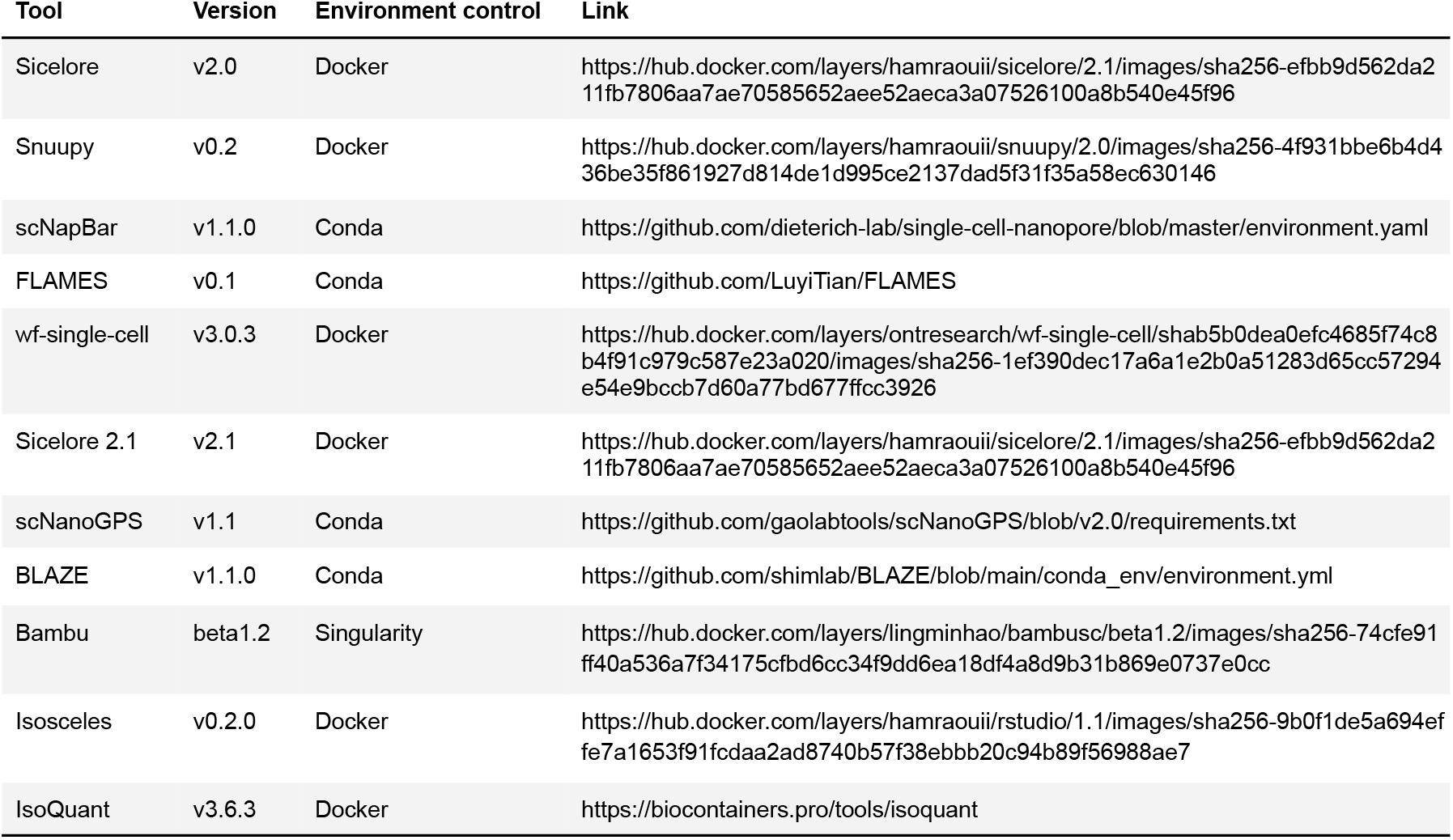
Software version and environment control strategies.

## Notes

https://github.com/alihamraoui/scKenver

https://zenodo.org/records/16098020

https://www.ebi.ac.uk/biostudies/arrayexpress/studies/E-MTAB-15190

### Competing Interest Statement

The authors have declared no competing interest.

## Bibliography

1. Amarinder Singh Thind, Isha Monga, Prasoon Kumar Thakur, Pallawi Kumari, Kiran Dindhoria, Monika Krzak, Marie Ranson, and Bruce Ashford. Demystifying emerging bulk RNA-Seq applications: the application and utility of bioinformatic methodology. Briefings in Bioinformatics, 22(6), nov 2021. doi: 10.1093/bib/bbab259.

2. Allon M. Klein, Linas Mazutis, Ilke Akartuna, Naren Tallapragada, Adrian Veres, Victor Li, Leonid Peshkin, David A. Weitz, and Marc W. Kirschner. Droplet Barcoding for Single-Cell Transcriptomics Applied to Embryonic Stem Cells. Cell, 161(5), May 2015. doi:10.1016/j.cell.2015.04.044.

3. Evan Z. Macosko, Anindita Basu, Rahul Satija, James Nemesh, Karthik Shekhar, Melissa Goldman, Itay Tirosh, Allison R. Bialas, Nolan Kamitaki, Emily M. Martersteck, John J. Trombetta, David A. Weitz, Joshua R. Sanes, Alex K. Shalek, Aviv Regev, and Steven A. McCarroll. Highly Parallel Genome-wide Expression Profiling of Individual Cells Using Nanoliter Droplets. Cell, 161(5), May 2015. doi: 10.1016/j.cell.2015.05.002.

4. Robert R. Stickels, Evan Murray, Pawan Kumar, Jilong Li, Jamie L. Marshall, Daniela J. Di Bella, Paola Arlotta, Evan Z. Macosko, and Fei Chen. Highly sensitive spatial transcriptomics at near-cellular resolution with Slide-seqV2. Nature Biotechnology, 39(3), March 2021. doi: 10.1038/s41587-020-0739-1.

5. Samuel G. Rodriques, Robert R. Stickels, Aleksandrina Goeva, Carly A. Martin, Evan Murray, Charles R. Vanderburg, Joshua Welch, Linlin M. Chen, Fei Chen, and Evan Z. Macosko. Slide-seq: A scalable technology for measuring genome-wide expression at high spatial resolution. Science, mar 2019. doi: 10.1126/science.aaw1219.

6. Christoph Ziegenhain, Beate Vieth, Swati Parekh, Björn Reinius, Amy Guillaumet-Adkins, Martha Smets, Heinrich Leonhardt, Holger Heyn, Ines Hellmann, and Wolfgang Enard. Comparative Analysis of Single-Cell RNA Sequencing Methods. Molecular Cell, 65(4), February 2017. doi: 10.1016/j.molcel.2017.01.023.

7. Roger Volden and Christopher Vollmers. Single-cell isoform analysis in human immune cells. Genome Biology, 23(1), December 2022. doi: 10.1186/s13059-022-02615-z.

8. Christoph Ziegenhain, Gert-Jan Hendriks, Michael Hagemann-Jensen, and Rickard Sandberg. Molecular spikes: a gold standard for single-cell RNA counting. Nature Methods, 19(5), January 2022. doi: 10.1038/s41592-022-01446-x.

9. Aziz M. Al’Khafaji, Jonathan T. Smith, Kiran V. Garimella, Mehrtash Babadi, Victoria Popic, Moshe Sade-Feldman, Michael Gatzen, Siranush Sarkizova, Marc A. Schwartz, Emily M. Blaum, Allyson Day, Maura Costello, Tera Bowers, Stacey Gabriel, Eric Banks, Anthony A. Philippakis, Genevieve M. Boland, Paul C. Blainey, and Nir Hacohen. High-throughput RNA isoform sequencing using programmed cDNA concatenation. Nature Biotechnology, 42(4), April 2024. doi: 10.1038/s41587-023-01815-7.

10. Ashley Byrne, Charles Cole, Roger Volden, and Christopher Vollmers. Realizing the potential of full-length transcriptome sequencing. Philosophical Transactions of the Royal Society B, nov 2019. doi: 10.1098/rstb.2019.0097.

11. Vincent Hahaut, Dinko Pavlinic, Walter Carbone, Sven Schuierer, Pierre Balmer, Mathieu Quinodoz, Magdalena Renner, Guglielmo Roma, Cameron S. Cowan, and Simone Picelli. Fast and highly sensitive full-length single-cell RNA sequencing using FLASH-seq. Nature Biotechnology, 40(10), October 2022. doi: 10.1038/s41587-022-01312-3.

12. Michael Hagemann-Jensen, Christoph Ziegenhain, and Rickard Sandberg. Scalable single-cell RNA sequencing from full transcripts with Smart-seq3xpress. Nature Biotechnology, 40 (10), October 2022. doi: 10.1038/s41587-022-01311-4.

13. Enze Deng, Qingmei Shen, Jingna Zhang, Yaowei Fang, Lei Chang, Guanzheng Luo, and Xiaoying Fan. Systematic evaluation of single-cell RNA-seq analyses performance based on long-read sequencing platforms. Journal of Advanced Research, May 2024. doi:10.1016/j.jare.2024.05.020.

14. Søren M. Karst, Ryan M. Ziels, Rasmus H. Kirkegaard, Emil A. Sørensen, Daniel McDonald, Qiyun Zhu, Rob Knight, and Mads Albertsen. High-accuracy long-read amplicon sequences 18 | bioRχiv Hamraoui et al. | Distributed under a Creative Commons Attribution | CC-BY 4.0 International license using unique molecular identifiers with Nanopore or PacBio sequencing. Nature Methods, 18(2), February 2021. doi: 10.1038/s41592-020-01041-y.

15. Carolina Monzó, Tianyuan Liu, and Ana Conesa. Transcriptomics in the era of long-read sequencing. Nature Reviews Genetics, mar 2025. doi: 10.1038/s41576-025-00828-z.

16. Kevin Lebrigand, Virginie Magnone, Pascal Barbry, and Rainer Waldmann. High throughput error corrected Nanopore single cell transcriptome sequencing. Nature Communications, 11(1), aug 2020. doi: 10.1038/s41467-020-17800-6.

17. Yanping Long, Zhijian Liu, Jinbu Jia, Weipeng Mo, Liang Fang, Dongdong Lu, Bo Liu, Hong Zhang, Wei Chen, and Jixian Zhai. FlsnRNA-seq: protoplasting-free full-length single-nucleus RNA profiling in plants. Genome Biology, 22(1), December 2021. doi: 10.1186/s13059-021-02288-0.

18. Qi Wang, Sven Bönigk, Volker Böhm, Niels Gehring, Janine Altmüller, and Christoph Dieterich. Single-cell transcriptome sequencing on the Nanopore platform with ScNapBar. RNA, 27(7), July 2021. doi: 10.1261/rna.078154.120.

19. Ghazal Ebrahimi, Baraa Orabi, Meghan Robinson, Cedric Chauve, Ryan Flannigan, and Faraz Hach. Fast and accurate matching of cellular barcodes across short-reads and long-reads of single-cell RNA-seq experiments. iScience, 25(7), jul 2022. doi: 10.1016/j.isci.2022.104530.

20. Cheng-Kai Shiau, Lina Lu, Rachel Kieser, Kazutaka Fukumura, Timothy Pan, Hsiao-Yun Lin, Jie Yang, Eric L. Tong, GaHyun Lee, Yuanqing Yan, Jason T. Huse, and Ruli Gao. High throughput single cell long-read sequencing analyses of same-cell genotypes and phenotypes in human tumors. Nature Communications, 14(1), jul 2023. doi: 10.1038/s41467-023-39813-7.

21. Yuntian Fu, Heonseok Kim, Sharmili Roy, Sijia Huang, Jenea I. Adams, Susan M. Grimes, Billy T. Lau, Anuja Sathe, Hanlee P. Ji, and Nancy R. Zhang. Single cell and spatial alternative splicing analysis with nanopore long read sequencing. Nature Communications, 16(6654), July 2025. doi: 10.1038/s41467-025-60902-2.

22. Peter De Rijk, Tijs Watzeels, Fahri Küçükali, Jasper Van Dongen, Júlia Faura, Patrick Willems, Lara De Deyn, Lena Duchateau, Carolin Grones, Thomas Eekhout, Tim De Pooter, Geert Joris, Stephane Rombauts, Bert De Rybel, Rosa Rademakers, Frank Van Breusegem, Mojca Strazisar, Kristel Sleegers, and Wouter De Coster. Scywalker: scalable end-to-end data analysis workflow for long-read single-cell transcriptome sequencing. Bioinformatics, 40(9), sep 2024. doi: 10.1093/bioinformatics/btae549.

23. Luyi Tian, Jafar S. Jabbari, Rachel Thijssen, Quentin Gouil, Shanika L. Amarasinghe, Oliver Voogd, Hasaru Kariyawasam, Mei R. M. Du, Jakob Schuster, Changqing Wang, Shian Su, Xueyi Dong, Charity W. Law, Alexis Lucattini, Yair David Joseph Prawer, Coralina Collar-Fernández, Jin D. Chung, Timur Naim, Audrey Chan, Chi Hai Ly, Gordon S. Lynch, James G. Ryall, Casey J. A. Anttila, Hongke Peng, Mary Ann Anderson, Christoffer Flensburg, Ian Majewski, Andrew W. Roberts, David C. S. Huang, Michael B. Clark, and Matthew E. Ritchie. Comprehensive characterization of single-cell full-length isoforms in human and mouse with long-read sequencing. Genome Biology, 22(1), nov 2021. doi: 10.1186/s13059-021-02525-6.

24. Andre Sim, Min Hao Ling, Ying Chen, Han Lu, Yi Xiang See, Arnaud Perrin, Ong Bee Leng Agnes, Elaine Yiqun Cao, Burton Chia, Jinyue Liu, Torsten Wüstefeld, Jay W. Shin, and Jonathan Göke. Isoform-level discovery, quantification and fusion analysis from single-cell and spatial long-read RNA-seq data with Bambu-Clump. bioRxiv, jan 2025. doi: 10.1101/2024.12.30.630828.

25. Yupei You, Yair D. J. Prawer, Ricardo De Paoli-Iseppi, Cameron P. J. Hunt, Clare L. Parish, Heejung Shim, and Michael B. Clark. Identification of cell barcodes from long-read single-cell RNA-seq with BLAZE. Genome Biology, 24(1), apr 2023. doi: 10.1186/s13059-023-02907-y.

26. Oliver Cheng, Min Hao Ling, Changqing Wang, Shuyi Wu, Matthew E. Ritchie, Jonathan Göke, Noorul Amin, and Nadia M. Davidson. Flexiplex: a versatile demultiplexer and search tool for omics data. Bioinformatics, 40(3), mar 2024. doi: 10.1093/bioinformatics/btae102.

27. Michal Kabza, Alexander Ritter, Ashley Byrne, Kostianna Sereti, Daniel Le, William Stephenson, and Timothy Sterne-Weiler. Accurate long-read transcript discovery and quantification at single-cell, pseudo-bulk and bulk resolution with Isosceles. Nature Communications, 15(1), aug 2024. doi: 10.1038/s41467-024-51584-3.

28. Andrey D. Prjibelski, Alla Mikheenko, Anoushka Joglekar, Alexander Smetanin, Julien Jarroux, Alla L. Lapidus, and Hagen U. Tilgner. Accurate isoform discovery with IsoQuant using long reads. Nature Biotechnology, 41(7), July 2023. doi: 10.1038/s41587-022-01565-y.

29. Björn E. Langer, Andreia Amaral, Marie-Odile Baudement, Franziska Bonath, Mathieu Charles, Praveen Krishna Chitneedi, Emily L. Clark, Paolo Di Tommaso, Sarah Djebali, Philip A. Ewels, Sonia Eynard, James A. Fellows Yates, Daniel Fischer, Evan W. Floden, Sylvain Foissac, Gisela Gabernet, Maxime U. Garcia, Gareth Gillard, Manu Kumar Gundappa, Cervin Guyomar, Christopher Hakkaart, Friederike Hanssen, Peter W. Harrison, Matthias Hörtenhuber, Cyril Kurylo, Christa Kühn, Sandrine Lagarrigue, Delphine Lallias, Daniel J. Macqueen, Edmund Miller, Júlia Mir-Pedrol, Gabriel Costa Monteiro Moreira, Sven Nahnsen, Harshil Patel, Alexander Peltzer, Frederique Pitel, Yuliaxis Ramayo-Caldas, Marcel da Câmara Ribeiro-Dantas, Dominique Rocha, Mazdak Salavati, Alexey Sokolov, Jose Espinosa-Carrasco, Cedric Notredame, and The Nf-Core Community. Empowering bioinformatics communities with Nextflow and nf-core. bioRxiv, may 2024. doi: 10.1101/2024.05.10.592912.

30. Philip A. Ewels, Alexander Peltzer, Sven Fillinger, Harshil Patel, Johannes Alneberg, Andreas Wilm, Maxime Ulysse Garcia, Paolo Di Tommaso, and Sven Nahnsen. The nf-core framework for community-curated bioinformatics pipelines. Nature Biotechnology, 38 (3), March 2020. doi: 10.1038/s41587-020-0439-x.

31. Austyn Trull, Nf-Core Community, Elizabeth A. Worthey, and Lara Ianov. scnanoseq: an nf-core pipeline for Oxford Nanopore single-cell RNA-sequencing. bioRxiv, apr 2025. doi: 10.1101/2025.04.08.647887.

32. Pallavi Gupta, Hannah O’Neill, Ernst J. Wolvetang, Aniruddha Chatterjee, and Ishaan Gupta. Advances in single-cell long-read sequencing technologies. NAR Genomics and Bioinformatics, 6(2), apr 2024. doi: 10.1093/nargab/lqae047.

33. Pallawi Kumari, Manmeet Kaur, Kiran Dindhoria, Bruce Ashford, Shanika L. Amarasinghe, and Amarinder Singh Thind. Advances in long-read single-cell transcriptomics. Human Genetics, 143(9), oct 2024. doi: 10.1007/s00439-024-02678-x.

34. Anne-Laure Boulesteix. Ten Simple Rules for Reducing Overoptimistic Reporting in Methodological Computational Research. PLOS Computational Biology, 11(4), apr 2015. doi: 10.1371/journal.pcbi.1004191.

35. Lukas M. Weber, Wouter Saelens, Robrecht Cannoodt, Charlotte Soneson, Alexander Hapfelmeier, Paul P. Gardner, Anne-Laure Boulesteix, Yvan Saeys, and Mark D. Robinson. Essential guidelines for computational method benchmarking. Genome Biology, 20(1), December 2019. doi: 10.1186/s13059-019-1738-8.

36. Salvador Capella-Gutierrez, Diana de la Iglesia, Juergen Haas, Analia Lourenco, José María Fernández, Dmitry Repchevsky, Christophe Dessimoz, Torsten Schwede, Cedric Notredame, Josep Ll Gelpi, and Alfonso Valencia. Lessons learned: Recommendations for establishing critical periodic scientific benchmarking. bioRxiv, 2017. doi: 10.1101/181677.

37. Serghei Mangul, Lana S. Martin, Brian L. Hill, Angela Ka-Mei Lam, Margaret G. Distler, Alex Zelikovsky, Eleazar Eskin, and Jonathan Flint. Systematic benchmarking of omics computational tools. Nature Communications, 10(1), mar 2019. doi: 10.1038/s41467-019-09406-4.

38. Xueyi Dong, Mei R. M. Du, Quentin Gouil, Luyi Tian, Jafar S. Jabbari, Rory Bowden, Pedro L. Baldoni, Yunshun Chen, Gordon K. Smyth, Shanika L. Amarasinghe, Charity W. Law, and Matthew E. Ritchie. Benchmarking long-read RNA-sequencing analysis tools using in silico mixtures. Nature Methods, 20(11), November 2023. doi: 10.1038/s41592-023-02026-3.

39. Yaqi Su, Zhejian Yu, Siqian Jin, Zhipeng Ai, Ruihong Yuan, Xinyi Chen, Ziwei Xue, Yixin Guo, D. Chen, Hongqing Liang, Zuozhu Liu, and Wanlu Liu. Comprehensive assessment of mRNA isoform detection methods for long-read sequencing data. Nature Communications, 15(1), may 2024. doi: 10.1038/s41467-024-48117-3.

40. Katarzyna J. Radomska, Audrey Onfroy, Laure Lecerf, Bastien Job, Aurélien Beaude, Laura Sesma Sanz, Tatiana El Jalkh, Denis Thieffry, Patrick Charnay, Pierre Wolkenstein, Nicolas Ortonne, Fanny Coulpier, and Piotr Topilko. Glial-to-mesenchymal transition of tumor schwann cells drives the genetic burden in mpnsts from neurofibromatosis type 1 mouse model. In Press, 2025.

41. Kevin Lebrigand, Joseph Bergenstråhle, Kim Thrane, Annelie Mollbrink, Konstantinos Meletis, Pascal Barbry, Rainer Waldmann, and Joakim Lundeberg. The spatial landscape of gene expression isoforms in tissue sections. Nucleic Acids Research, 51(8), may 2023. doi: 10.1093/nar/gkad169.

42. Ali Hamraoui, Laurent Jourdren, and Morgane Thomas-Chollier. AsaruSim: a single-cell and spatial RNA-Seq Nanopore long-reads simulation workflow. Bioinformatics, feb 2025. doi: 10.1093/bioinformatics/btaf087.

43. Tom Smith, Andreas Heger, and Ian Sudbery. UMI-tools: modeling sequencing errors in Unique Molecular Identifiers to improve quantification accuracy. Genome Research, 27(3), March 2017. doi: 10.1101/gr.209601.116.

44. Dirk Merkel. Docker: lightweight linux containers for consistent development and deployment. Linux journal, 2014(239), 2014. doi: 10.5555/2600239.2600241.

45. Aaron T. L. Lun, Samantha Riesenfeld, Tallulah Andrews, The Phuong Dao, Tomas Gomes, participants in the 1st Human Cell Atlas Jamboree, and John C. Marioni. EmptyDrops: distinguishing cells from empty droplets in droplet-based single-cell RNA sequencing data. Genome Biology, 20(1), December 2019. doi: 10.1186/s13059-019-1662-y.

46. Ilya Korsunsky, Nghia Millard, Jean Fan, Kamil Slowikowski, Fan Zhang, Kevin Wei, Yuriy Baglaenko, Michael Brenner, Po-ru Loh, and Soumya Raychaudhuri. Fast, sensitive and accurate integration of single-cell data with Harmony. Nature Methods, 16(12), December 2019. doi: 10.1038/s41592-019-0619-0.

47. Lulu Shang and Xiang Zhou. Spatially aware dimension reduction for spatial transcriptomics. Nature Communications, 13(1), nov 2022. doi: 10.1038/s41467-022-34879-1.

48. Zhijian Li, Zain M. Patel, Dongyuan Song, Guanao Yan, Jingyi Jessica Li, and Luca Pinello. Benchmarking computational methods to identify spatially variable genes and peaks. bioRxiv, dec 2023. doi: 10.1101/2023.12.02.569717.

49. Eline Beert, Hilde Brems, Bruno Daniëls, Ivo De Wever, Frank Van Calenbergh, Joseph Schoenaers, Maria Debiec-Rychter, Olivier Gevaert, Thomas De Raedt, Annick Van Den Bruel, Thomy Ravel, Karen Cichowski, Lan Kluwe, Victor Mautner, Raf Sciot, and Eric Legius. Atypical neurofibromas in neurofibromatosis type 1 are premalignant tumors. Genes, Chromosomes and Cancer, 50(12), dec 2011. doi: 10.1002/gcc.20921.

50. Shanrong Zhao, Alexander Barron, and Ken Dower. Rna Sequencing to Explore Dominant Isoform Switch During CCR6+ Memory T Cell Activation. Preprint on Research Square, jan 2021. doi: 10.21203/rs.3.rs-140817/v1.

51. Xiaoxia Che, Xin Guan, Yiyin Ruan, Lifei Shen, Yuhong Shen, Hua Liu, Chongying Zhu, Tianyu Zhou, Yiwei Wang, and Weiwei Feng. TRIM4 modulates the ubiquitin-mediated degradation of hnRNPDL and weakens sensitivity to CDK4/6 inhibitor in ovarian cancer. Frontiers of Medicine, 19(1), feb 2025. doi: 10.1007/s11684-024-1103-5.

52. Geo Pertea and Mihaela Pertea. Gff Utilities: Gffread and GffCompare. F1000Research, 9, sep 2020. doi: 10.12688/f1000research.23297.2.

53. Elisabetta Mereu, Atefeh Lafzi, Catia Moutinho, Christoph Ziegenhain, Davis J. McCarthy, Adrián Álvarez-Varela, Eduard Batlle Sagar, Dominic Grün, Julia K. Lau, Stéphane C. Boutet, Chad Sanada, Aik Ooi, Robert C. Jones, Kelly Kaihara, Chris Brampton, Yasha Talaga, Yohei Sasagawa, Kaori Tanaka, Tetsutaro Hayashi, Caroline Braeuning, Cornelius Fischer, Sascha Sauer, Timo Trefzer, Christian Conrad, Xian Adiconis, Lan T. Nguyen, Aviv Regev, Joshua Z. Levin, Swati Parekh, Aleksandar Janjic, Lucas E. Wange, Johannes W. Bagnoli, Wolfgang Enard, Marta Gut, Rickard Sandberg, Itoshi Nikaido, Ivo Gut, Oliver Stegle, and Holger Heyn. Benchmarking single-cell RNA-sequencing protocols for cell atlas projects. Nature Biotechnology, 38(6), June 2020. doi: 10.1038/s41587-020-0469-4.

54. Arthur Dondi, Ulrike Lischetti, Francis Jacob, Franziska Singer, Nico Borgsmüller, Ricardo Coelho, Viola Heinzelmann-Schwarz, Christian Beisel, and Niko Beerenwinkel. Detection of isoforms and genomic alterations by high-throughput full-length single-cell RNA sequencing in ovarian cancer. Nature Communications, 14(1), nov 2023. doi: 10.1038/s41467-023-43387-9.

55. Francisco J. Pardo-Palacios, Dingjie Wang, Fairlie Reese, Mark Diekhans, Sílvia Carbonell-Sala, Brian Williams, Jane E. Loveland, Maite D. María, Matthew S. Adams, Gabriela Balderrama-Gutierrez, Amit K. Behera, Jose M. Gonzalez Martinez, Toby Hunt, Julien Lagarde, Cindy E. Liang, Haoran Li, Marcus Jerryd Meade, David A. Moraga Amador, Andrey D. Prjibelski, Inanc Birol, Hamed Bostan, Ashley M. Brooks, Muhammed Hasan Çelik, Ying Chen, Mei R. M. Du, Colette Felton, Jonathan Göke, Saber Hafezqorani, Ralf Herwig, Hideya Kawaji, Joseph Lee, Jian-Liang Li, Matthias Lienhard, Alla Mikheenko, Dennis Mulligan, Ka Ming Nip, Mihaela Pertea, Matthew E. Ritchie, Andre D. Sim, Alison D. Tang, Yuk Kei Wan, Changqing Wang, Brandon Y. Wong, Chen Yang, If Barnes, Andrew E. Berry, Salvador Capella-Gutierrez, Alyssa Cousineau, Namrita Dhillon, Jose M. Fernandez-Gonzalez, Luis Ferrández-Peral, Natàlia Garcia-Reyero, Stefan Götz, Carles Hernández-Ferrer, Liudmyla Kondratova, Tianyuan Liu, Alessandra Martinez-Martin, Carlos Menor, Jorge Mestre-Tomás, Jonathan M. Mudge, Nedka G. Panayotova, Alejandro Pafniagua, Dmitry Repchevsky, Xingjie Ren, Eric Rouchka, Brandon Saint-John, Enrique Sapena, Leon Sheynkman, Melissa Laird Smith, Marie-Marthe Suner, Hazuki Takahashi, Ingrid A. Youngworth, Piero Carninci, Nancy D. Denslow, Roderic Guigó, Margaret E. Hunter, Rene Maehr, Yin Shen, Hagen U. Tilgner, Barbara J. Wold, Christopher Vollmers, Adam Frankish, Kin Fai Au, Gloria M. Sheynkman, Ali Mortazavi, Ana Conesa, and Angela N. Brooks. Systematic assessment of long-read RNA-seq methods for transcript identification and quantification. Nature Methods, 21(7), July 2024. doi: 10.1038/s41592-024-02298-3.

56. Leandro Lima, Camille Marchet, Ségolène Caboche, Corinne Da Silva, Benjamin Istace, Jean-Marc Aury, Hélène Touzet, and Rayan Chikhi. Comparative assessment of long-read error correction software applied to nanopore rna-sequencing data. Briefings in Bioinformatics, 21(4), 06 2019. doi: 10.1093/bib/bbz058.

57. Ying Chen, Nadia M. Davidson, Yuk Kei Wan, Fei Yao, Yan Su, Hasindu Gamaarachchi, Andre Sim, Harshil Patel, Hwee Meng Low, Christopher Hendra, Laura Wratten, Christopher Hakkaart, Chelsea Sawyer, Viktoriia Iakovleva, Puay Leng Lee, Lixia Xin, Hui En Vanessa Ng, Jia Min Loo, Xuewen Ong, Hui Qi Amanda Ng, Jiaxu Wang, Wei Qian Casslynn Koh, Suk Yeah Polly Poon, Dominik Stanojevic, Hoang-Dai Tran, Kok Hao Edwin Lim, Shen Yon Toh, Philip Andrew Ewels, Huck-Hui Ng, N. Gopalakrishna Iyer, Alexandre Thiery, Wee Joo Chng, Leilei Chen, Ramanuj DasGupta, Mile Sikic, Yun-Shen Chan, Boon Ooi Patrick Tan, Yue Wan, Wai Leong Tam, Qiang Yu, Chiea Chuan Khor, Torsten Wüstefeld, Alexander Lezhava, Ploy N. Pratanwanich, Michael I. Love, Wee Siong Sho Goh, Sarah B. Ng, Alicia Oshlack, SG-NEx consortium, N. Gopalakrishna Iyer, Qiang Yu, and Jonathan Göke. A systematic benchmark of Nanopore long-read RNA sequencing for transcript-level analysis in human cell lines. Nature Methods, mar 2025. doi: 10.1038/s41592-025-02623-4.

58. Hoa Thi Nhu Tran, Kok Siong Ang, Marion Chevrier, Xiaomeng Zhang, Nicole Yee Shin Lee, Michelle Goh, and Jinmiao Chen. A benchmark of batch-effect correction methods for single-cell RNA sequencing data. Genome Biology, 21(1), December 2020. doi: 10.1186/s13059-019-1850-9.

59. Malte D. Luecken, M. Büttner, K. Chaichoompu, A. Danese, M. Interlandi, M. F. Mueller, D. C. Strobl, L. Zappia, M. Dugas, M. Colomé-Tatché, and Fabian J. Theis. Benchmarking atlas-level data integration in single-cell genomics. Nature Methods, 19(1), January 2022. doi: 10.1038/s41592-021-01336-8.

60. Xin Chen, Zhaowei Yang, Wanqiu Chen, Yongmei Zhao, Andrew Farmer, Bao Tran, Vyacheslav Furtak, Malcolm Moos, Wenming Xiao, and Charles Wang. A multi-center cross-platform single-cell RNA sequencing reference dataset. Scientific Data, 8(1), feb 2021. doi: 10.1038/s41597-021-00809-x.

61. Heng Li. Minimap2: pairwise alignment for nucleotide sequences. Bioinformatics, 34(18), sep 2018. doi: 10.1093/bioinformatics/bty191.

62. Li, H. Github-lh3/seqtk: toolkit for processing sequences in FASTA/Q formats, 2021.

63. Jacques Dainat. Agat: Another gff analysis toolkit to handle annotations in any gtf/gff format, 2022.

64. Pierre Lindenbaum and Richard Redon. bioalcidae, samjs and vcffilterjs: object-oriented formatters and filters for bioinformatics files. Bioinformatics, 34(7), November 2017. doi: 10.1093/bioinformatics/btx734.

65. Yuhan Hao, Stephanie Hao, Erica Andersen-Nissen, William M. Mauck, Shiwei Zheng, Andrew Butler, Maddie J. Lee, Aaron J. Wilk, Charlotte Darby, Michael Zager, Paul Hoffman, Marlon Stoeckius, Efthymia Papalexi, Eleni P. Mimitou, Jaison Jain, Avi Srivastava, Tim Stuart, Lamar M. Fleming, Bertrand Yeung, Angela J. Rogers, Juliana M. McElrath, Catherine A. Blish, Raphael Gottardo, Peter Smibert, and Rahul Satija. Integrated analysis of multimodal single-cell data. Cell, 184(13), June 2021. doi: 10.1016/j.cell.2021.04.048.

66. Lei Guo, Zhenxing Hu, Chao Zhao, Xiangnan Xu, Shujuan Wang, Jingjing Xu, Jiyang Dong, and Zongwei Cai. Data Filtering and Its Prioritization in Pipelines for Spatial Segmentation of Mass Spectrometry Imaging. Analytical Chemistry, mar 2021. doi: 10.1021/acs.analchem.0c05242.

67. Theodore Alexandrov and Andreas Bartels. Testing for presence of known and unknown molecules in imaging mass spectrometry. Bioinformatics, 29(18), sep 2013. doi: 10.1093/bioinformatics/btt388.

68. Guangchuang Yu, Li-Gen Wang, Yanyan Han, and Qing-Yu He. clusterProfiler: an R Package for Comparing Biological Themes Among Gene Clusters. OMICS: A Journal of Integrative Biology, 16(5), May 2012. doi: 10.1089/omi.2011.0118.

69. Yu Hu, Li Fang, Xuelian Chen, Jiang F. Zhong, Mingyao Li, and Kai Wang. LIQA: long-read isoform quantification and analysis. Genome Biology, 22(1), December 2021. doi: 10.1186/s13059-021-02399-8.

70. Manuel Tardaguila, Lorena Fuente, Cristina Marti, Cécile Pereira, Francisco Jose Pardo-Palacios, Hector Risco, Marc Ferrell, Maravillas Mellado, Marissa Macchietto, Kenneth Verheggen, Mariola Edelmann, Iakes Ezkurdia, Jesus Vazquez, Michael Tress, Ali Mortazavi, Lennart Martens, Susana Rodriguez-Navarro, Victoria Moreno-Manzano, and Ana Conesa. Sqanti: extensive characterization of long-read transcript sequences for quality control in full-length transcriptome identification and quantification. Genome Research, 28(3), jan 2018. doi: 10.1101/gr.222976.117.

